# Graph models of brain state in deep anaesthesia reveal sink state dynamics of reduced spatiotemporal complexity

**DOI:** 10.1101/2024.08.21.608935

**Authors:** James Wilsenach, Charlotte M. Deane, Gesine Reinert, Katie Warnaby

**Affiliations:** Alan Turing Institute, London, United Kingdom; Department of Statistics, University of Oxford, Oxford, United Kingdom; Oxford Centre for Functional MRI of the Brain, University of Oxford, Oxford, United Kingdom

**Keywords:** dynamic brain state models, consciousness, complexity, spatiotemporal community detection, Hidden Markov models, anaesthesia

## Abstract

Anaesthetisia is an important surgical and explorative tool in the study of consciousness. Much work has been done to connect the deeply anaesthetised condition with decreased complexity. However, anaesthesia-induced unconsciousness is also a dynamic condition in which functional activity and complexity may fluctuate, being perturbed by internal or external (e.g. noxious) stimuli. We use fMRI data from a cohort undergoing deep propofol anaesthesia to investigate resting state dynamics using dynamic brain state models and spatiotemporal network analysis. We focus our analysis on group-level dynamics of brain state temporal complexity, functional activity, connectivity and spatiotemporal modularization in deep anaesthesia and wakefulness. We find that in contrast to dynamics in the wakeful condition, anaesthesia dynamics are dominated by a handful of sink states that act as low-complexity attractors to which subjects repeatedly return. On a subject-level, our analysis provides tentative evidence that these low complexity attractor states appear to depend on subject-specific age and anaesthesia susceptibility factors. Finally, our spatiotemporal analysis, including a novel spatiotemporal clustering of graphs representing hidden Markov models, suggests that dynamic functional organisation in anaesthesia can be characterised by mostly unchanging, isolated regional subnetworks that share some similarities with the brain’s underlying structural connectivity, as determined from normative tractography data.

## INTRODUCTION

General anaesthesia is an invaluable surgical tool that reduces the overall risk of complications and shields patients from immediate pain while being generally safe and effective Urban and Bleckwenn (2002). However, anaesthetic susceptibility and risks are patient specific and may include impaired respiration and heart function Nunn (1990), delayed recovery of consciousness and post-operative delirium in over-anaesthesia Bryson and Wyand (2006); Fritz et al. (2016). Over-anaesthetisation has been associated with cognitive decline in older patients who are also at higher risk of post-operative delirium Sevenius Nilsen, Juel, and Marshall (2019); Wu, Zhao, Weng, and Ma (2019). Conversely, under-anaesthetisation may risk psychological trauma through post-surgical recollection of pain and helplessness, especially in combination with commonly administered muscle relaxants Aceto et al. (2013); Schwender et al. (1998); Urban and Bleckwenn (2002). These factors motivate the importance of understanding how the brain dynamically maintains unconsciousness under sustained anaesthesia when subjected to passive, or, in the case of surgery, powerful forms of external stimulation Bonhomme et al. (2019).

More generally, understanding dynamic functional changes in neural configuration, or *brain state*, has been key to describing the processes involved in the transitions between and maintenance of different conscious and unconsciousness conditions He, Das, Hotan, and Purdon (2023); Mediano et al. (2023); C. Zhang et al. (2024). A popular method for determination of brain state space models has been **Hidden Markov Models (HMMs)**, with applications to sleep Stevner et al. (2019); Yang et al. (2024), to Disorders of Consciousness (DOCs) Bai et al. (2021), and to anaesthesia Liu et al. (2022). In HMMs, multiple signals of brain activity are generally modelled as arising from discrete hidden states; see Supplementary Section S.1 for a brief introduction. These states provide a snapshot of functional activity and connectivity that subjects transition between, probabilistically and dynamically, to support the maintenance of a condition such as aneasthesia-induced unconsciousness or resting wakefulness Vidaurre et al. (2016). HMMs are appealing in part because of this rich spatiotemporal representations of dynamics that depend on underlying persistent and stable conditions such as in a resting state Leroux (1992). Stability dependence differentiates HMMs from more sophisticated models of brain state which may be applied to non-stationary behaviours, usually at the cost of the richness of temporal dynamics in the model Ito, Yang, Laurent, Schultz, and Cole (2022); Parks et al. (2024); Rukat, Baker, Quinn, and Woolrich (2016). We focus our study on HMM analysis of resting brain state dynamics in both wakefulness and anaesthesia-induced unconsciousness in order to better understand the dynamic properties that support its stability and maintenance.

HMMs and other state-space models have been used to demonstrate a general decrease in temporal Demertzi et al. (2019); Luppi et al. (2024); Stevner et al. (2019) and signal-based complexity measures Liang et al. (2024); Ruffini (2017) during unconsciousness including under most anaesthetics, with the possible exception of ketamine Li and Mashour (2019). Of these measures, Kolmogorov complexity is thought to be a possible neural correlate of consciousness and is notable for its strong links to other proposed neural correlates such as in Integrated Information and the Global Workspace Fan, Yeh, Chen, Shieh, et al. (2011); Marshall, Gomez-Ramirez, and Tononi (2016); Ruffini (2017), though its accurate quantification in practice in particular in recordings of brain activity remains computationally and mathematically intractable Sevenius Nilsen et al. (2019). Using state-space models however, the Kolmogorov complexity can be approximated by the *entropy* or *information rate* Galatolo, Hoyrup, and Rojas (2010). This approximation has been used to demonstrate a general decrease in the dynamic reachability or realisability of states during various forms of unconsciousness when compared to wakefulness Barttfeld et al. (2015); Castro et al. (2024); Demertzi et al. (2019); Li, Vlisides, and Mashour (2022); Mediano et al. (2023). We corroborate these findings by training HMMs which are specific to each condition, wakefulness and deep anaesthesia, and propose that certain condition-specific states may act as low-complexity attractors or *sinks* which are disproportionately responsible for this reduction in complexity.

Brain states can be considered as graphs connecting brain regions, with edges depending on the functional connectivity between these regions Vidaurre et al. (2016). This representation has helped to establish the minimal resting state network features which persist to support unconsciousness dynamics Barttfeld et al. (2015); Heine et al. (2012); Luppi et al. (2019). However, consideration of states in isolation fails to incorporate the temporal dependencies in the activity of these subnetworks. Therefore, in analogy to the static spatially-defined resting state networks Heine et al. (2012). Here we provide a spatiotemporal framework for brain state analysis to understand the spatiotemporal functional modularisation of activity in both conditions, wakefulness and anaesthesia. To this purpose we represent a fitted HMM on brain states as a graph, with brain states as nodes, and the transition probabilities between brain states taken as edge weights between nodes; the resulting graph on graphs is a **Hidden Markov Graph Model (HMGM)** as in Wilsenach, Warnaby, Deane, and Reinert (2022). This particular graph on graphs structure allows us to develop new methods to detect modular spatiotemporal subnetworks or **communities**, incorporating both functional and temporal information. In particular, we detect spatiotemporal functional stratification which occurs in anaesthesia-induced unconsciousness but not wakefulness. We also use these methods to corroborate and expand upon previous work on the close links between structure and unconscious brain state connectivity Barttfeld et al. (2015); López-González et al. (2021); Luppi et al. (2023).

Our results suggest new avenues for exploration in the study of anaesthesia, in particular how age and anaesthesia susceptibility may modulate patient trajectories both into and out of anaesthesia-induced unconsciousness. Moreover, we demonstrate that unconsciousness dynamics may involve effective restrictions on the space of possible brain states through dominant *sink* states. We find indications of the dissolution of important **functional modules** and the loss of the brain’s capacity for integrating multiple streams of information under **propofol**-induced deep anaesthesia.

## MATERIALS AND METHODS

Fig. 1 shows an overview of the experimental set-up (Panel A) and the processing of the blood oxygen level-dependent (**BOLD**) signals, followed by training of the HMM (Panel B); Panel C shows the construction of the HMGM from the trained HMM, and how the brain **Regions of Interest (ROIs)** and states can be decomposed in terms of both their functional module or community memberships. This framework builds upon previous work first introduced in Wilsenach (2022) and Wilsenach et al. (2022).

**Figure 1:**
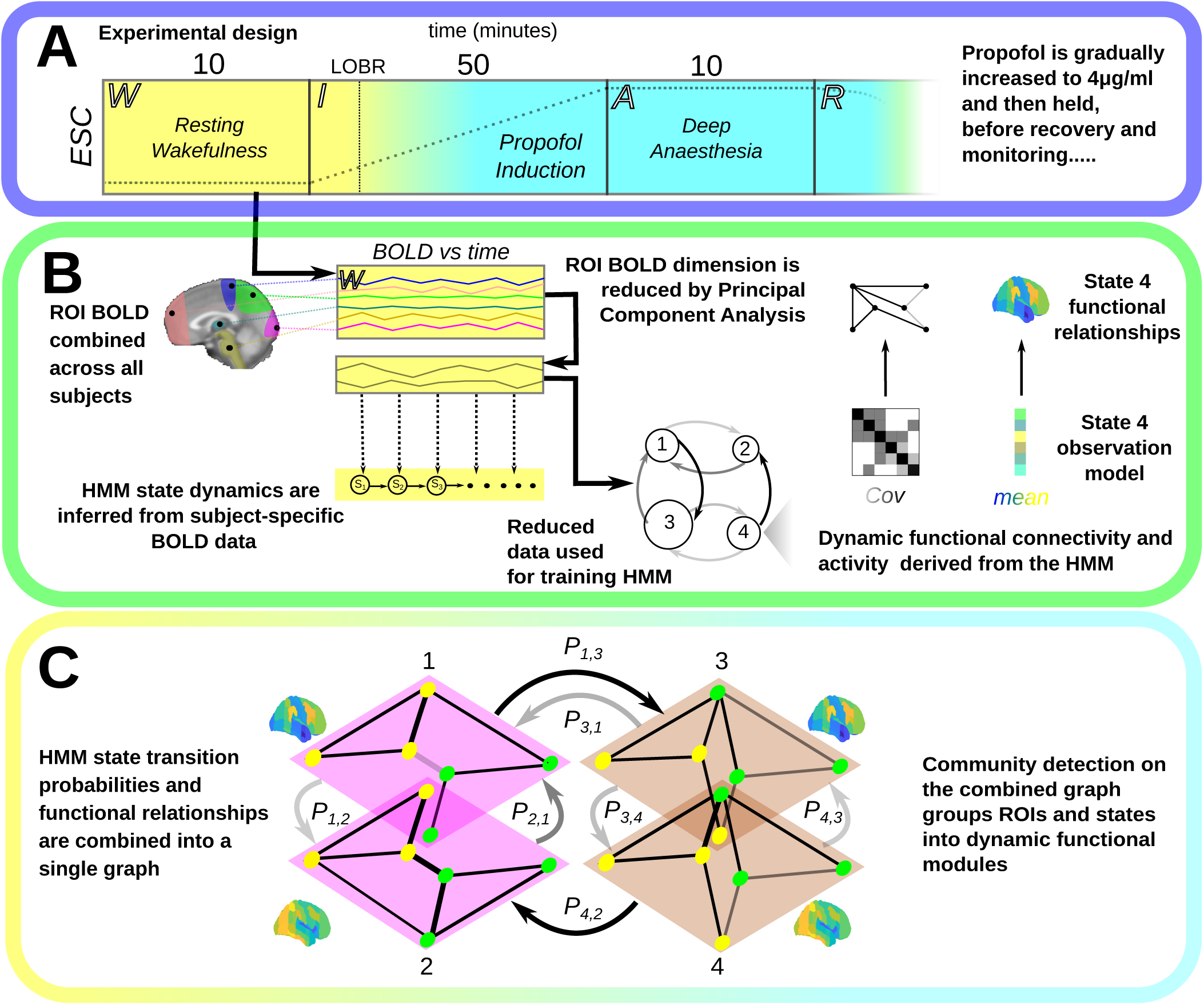
This figure shows our experimental and analysis framework. (A) Subject fMRI is recorded at resting wakefulness, yellow, W, followed by ultra-slow propofol induction, I, where the estimated Effect Site Concentration (ESC) is gradually increased, and **Loss of Behavioural Responsiveness (LOBR)** during induction is marked with a dotted line, before being held at 4*µ*g/ml during the deep anaesthesia, cyan, phase A. Anaesthesia dosage is then tapered and subjects are allowed to recover, R. (B) ROI BOLD time series across all subjects were first combined and then dimensionality reduced by Principal Component Analysis (PCA). In the framework, brain dynamics arise from a sequence of hidden brain states. Occurrence of the next state is probabilistically dependent on the current state. These probabilities are shown with an arrow (darker, higher probability). Each state encodes expected functional connectivity and activity relationships that are mapped into parcellation space through the state mean (blue, low) and covariance (dark, high) parameters. The analysis is performed again for anaesthesia. (D) We combine functional and dynamic information to produce the combined HMGM graph (dark arrows, high probability transition), followed by spatiotemporal community detection.

### Data Acquisition and fMRI Pre-processing

BOLD fMRI data was recorded over the course of a four phase experiment in which 16 initial subjects received a phase-specific dose of the GABA-ergic general anaesthetic propofol Mhuircheartaigh, Warnaby, Rogers, Jbabdi, and Tracey (2013). In the first phase, baseline wakeful activity was recorded for 10 minutes. Subjects were asked to remain still with eyes closed. In the second phase, propofol was gradually infused to a maximum **Effect Site Concentration (ESC)** of 4*µg/ml*, which is well above the minimal dose previously found to be effective in inducing loss of consciousness in subjects and is sufficient for inducing sleep-like slow waves Forrest, Tooley, Saunders, and Prys-Roberts (1994). In phase three the maximum ESC was maintained for 10 minutes, leading to a stable deep anaesthesia state. In phase four, the propofol dose was gradually decreased until subjects regained full responsiveness. The experimental overview is shown in Fig. 1A; Supplementary Section S.2 contains greater detail.

The frequency of the fMRI data was one full head volume every three seconds at a 3D voxel resolution of 2mm^3^. All brain volumes were head extracted, artefact corrected and registered to a single 1mm^3^, subject-specific high-resolution volume-scan in FSL Jenkinson, Beckmann, Behrens, Woolrich, and Smith (2012). The resulting volumes were mapped into the MNI152 MRI standard space. To obtain a computationally tractable anatomical partition with cortical-subcortical coverage, we used the Harvard-Oxford (HO) cortical and sub-cortical atlases with *D* = 63 ROIs developed in Desikan et al. (2006) and made into a combined atlas in Wilsenach et al. (2022).

In this analysis we focus on the 10 minutes pre-induction (phase 1) and the 10 minutes post-induction recordings (phase 3), when subjects are held at the maximum dosage. The high dosage results in a condition of deep anaesthesia-induced unconsciousness and non-responsiveness to conventional rousing stimuli. Subjects were excluded based-on whether they exhibited EEG signatures that have been linked with poor tolerance for high anaesthesia dosages and post-operative delirium (see Supplementary Section S.3) Fritz et al. (2016); Z. Zhang et al. (2019). After exclusion the total number of complete recordings for subjects in both conditions was *N* = 13.

In order to study the relationship of the brain states to the brain’s structural network we accessed a large structural dataset combining data from 169 normative adult **tractographies** based on multi-shell Diffusion Tensor Imaging re-registered to MNI152 standard space. The tractographies were constructed using the Gibbs tracking algorithm Reisert et al. (2011), which were then co-registered in standard space and combined as outlined in Horn and Blankenburg (2016).

### Model Training and State Selection

HMM model selection and training for both anaesthesia and wakefulness was carried out as detailed in Wilsenach et al. (2022), using the Variational Bayes HMM training procedure in Vidaurre et al. (2016). This procedure is shown in Fig. 1B for wakefulness (and is carried out analogously in anaesthesia). As the data are high-dimensional, some compression is needed for computational efficiency. First the high resolution voxel-wise data is reduced by obtaining the mean BOLD signal across each ROI over the recording. For each ROI, the recordings for all subjects are then concatenated. To further reduce computational and model complexity in the HMM fitting process, a Principal Component Analysis was carried out to find computationally efficient combinations of ROI series, with the reduced dimension selected by parallel analysis (see Supplementary Section S.4). The reduced data was then used to train an HMM. The observations of the HMM are thus not ROIs themselves but instead combinations of ROIs. As the concatenated data obstructs the subject separation, subject-specific time series are obtained from the PCA-reduced data to infer state probabilities at each time point, while respecting the separation between subject recordings.

Following HMM training, the number of initial HMM states was selected by maximum entropy. The maximum entropy method has been shown to broadly agree with other methods of model size selection such as cross-validated maximum likelihood Wilsenach (2022), while often resulting in larger models that allow for more complex possible dynamics Wilsenach et al. (2022). As this selection may result in states that are not robustly expressed, HMM states were removed if they appeared in less than 20% of subjects, and transition probabilities were re-standardised so that each row sums to one (see Supplementary Fig. 3). The (fitted) probability of each HMM state occurrence at each time point is then calculated, which determines the **Fractional Occupancy (FO)** of each state. This is the expected fraction of time spent in each state over the length of a subject recording. The resulting model for each condition has a total of *K_wake_* or *K_anaes_* states respectively and *D* ROIs.

### Graph Models of Dynamic Function and Structure

HMGMs are spatiotemporal graph models derived from the respective HMMs. More formally, an HMGM is a **multiplex** graph with each layer representing a brain state with the same set of ROIs as nodes, having edges determined by the functional connectivity between the ROIs, and in which the interlayer edges are the probabilities of transition between brain states. This structure can be seen in Fig. 1C. HMGMs were derived from the HMMs for anaesthesia and wakefulness respectively.

A simpler, single-layer structural graph model was fit to the tractography data using DSI-studio’s structural connectome tool Yeh, Verstynen, Wang, Fernández-Miranda, and Tseng (2013) and the HO atlas with default parameters and median streamline length normalisation, producing an undirected, weighted graph model proxy of subject structural connectivity Yeh et al. (2018).

### Consensus Community Detection

One of the primary strengths of graph-based models is that they allow for the decomposition of the graph into functionally related modules or communities. Modularity-based community detection is a collection of methods for finding optimal groupings of nodes (brain regions) in a graph so that members of the group are preferentially connected to one another Girvan and Newman (2002); Newman (2006) in a way which numerically aims to maximise the *modularity* of the partition.

In order to determine the underlying *structural* modular organisation of the structural graph model we applied a robust version of the modularity optimisation method based on Lancichinetti and Fortunato (2012) using multiple parameters and initialisations to determine structural communities (see Supplementary Section S.6 for further details).

### Beyond Static Communities…

Spatiotemporal communities can be interpreted as groups of brain regions that tend to coordinate not only in space but also over time. A generalised form of modularity-based community detection was used to partition the nodes (ROIs) of the HMGM into functionally related sets that coordinate activity across space and time into dynamic functional modules (node colour in Fig. 1C). We determine the spatiotemporal community structure of each of the HMGM models using a new modification to the modularity that combines spatial and temporal HMM properties. The modified score takes into account both functional connectivity and temporal probabilities of transition between states Mucha, Richardson, Macon, Porter, and Onnela (2010); Sen, Chu, and Parhi (2019). Using this score, communities can be interpreted as representing functional modules which can reorganise across time in order to react to external and internal stimuli. In all cases the modularity is optimised using the generalised Louvain algorithm Mucha et al. (2010), with the same robust aggregation over multiple parameters as in Lancichinetti and Fortunato (2012); see Supplementary Section S.6 for more detail on these methods and justification of the range of parameters. Layers with similar community composition are also clustered hierarchically; in Fig. 1C members of the same cluster are given the same colour.

### Sink States, Switching and Information Rates

HMMs are probabilistic state space models in which state transitions occur probabilistically depending only on the previous state in the sequence. In a sufficiently regular Markov model, the stationary probability of a state is the long-term probability of occupying this state over a sufficiently long time (see Supplementary Section S.7). Here, we use the stationary probability to define a centrality measure over states. In analogy to PageRank centrality, we term the stationary probability of each state its *sink centrality*. As our interest lies in those states which cumulatively account for the majority of the expected time spent, in each condition, to simplify the analysis, special focus was then put on “strong” sink states that disproportionately account for activity under each conditions. Here we use as cut-off for a sink state to be considered ‘strong’ that it has a sink centrality of at least 0.05. Other threshold choices are possible; the threshold is chosen purely to summarise the model behaviour, not for training the model and is explored further in Results.

Fig. 2 provides a sketch of how HMM and HMGM analysis can provide important axes along which to examine brain state activity by contrasting model behaviours. The switching rate, Fig. 2A, is a well established measure of the temporal complexity of brain state models Stevner et al. (2019); Vidaurre et al. (2016). It is the frequency at which state change occurs in the state trajectory that best fits the data (see Supplementary Section S.8) Brin and Page (1998). Fig. 2A shows four different four-state Markov models, where node size is proportional to the sink centrality. Edges are shaded from high (dark) to low (light) probability. A plausible state sequence is generated from each model from which the state switching events are determined and the switching rate (in Hz) is calculated. The FO is also shown, which is based on the expected time spent in each state, which is inferred directly from the model (see Supplementary Section S.1). The FO can be viewed as the distribution describing the probability of observing a subject in a given state during an entire recording.

**Figure 2:**
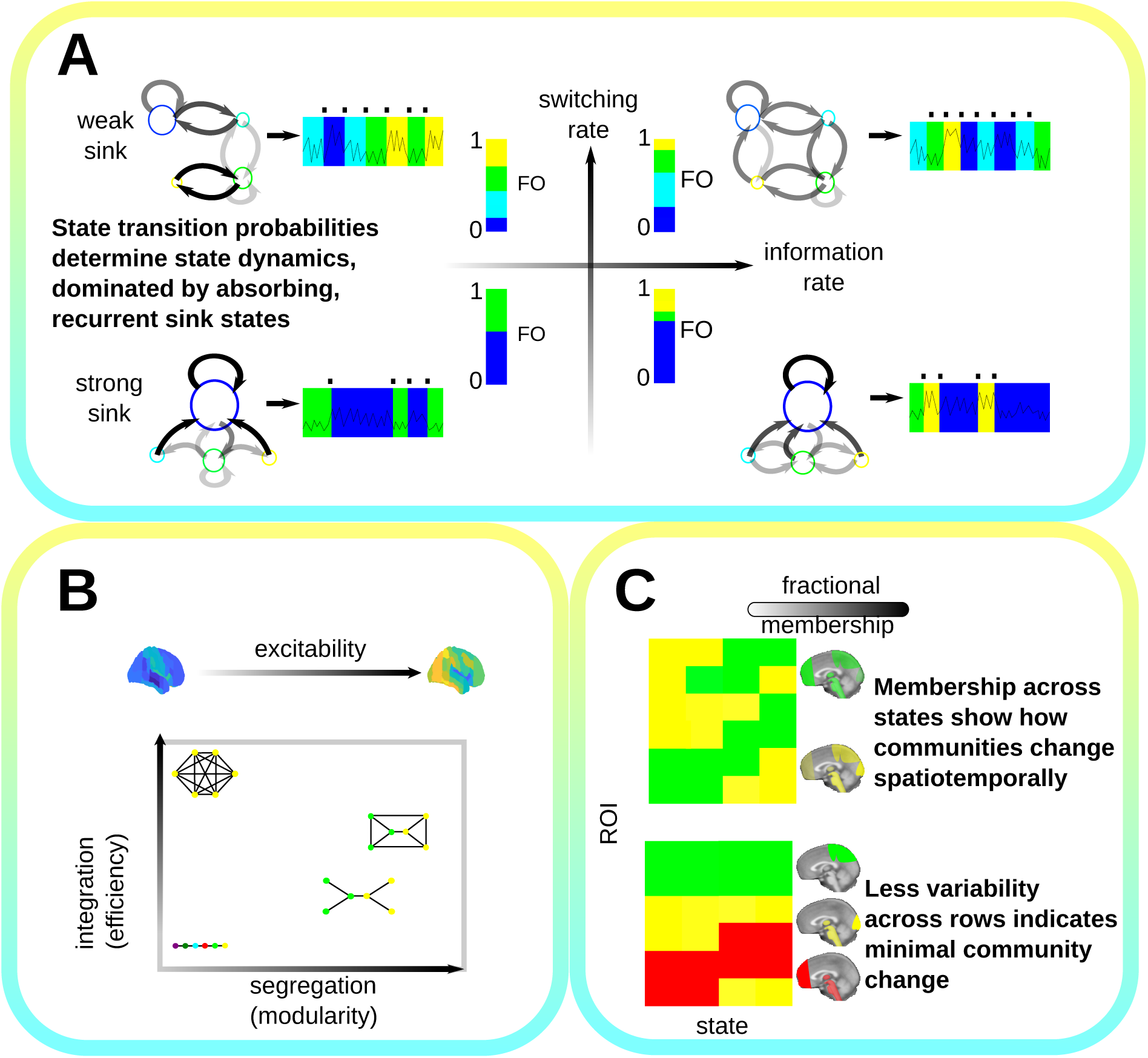
This figure shows our analysis framework (A) Four state models are shown to contrast their information theoretic properties along two axes switching rate increasing from top to bottom and information (entropy) rate increasing from left to right. Larger nodes correspond to sinks. Fractional Occupancy (FO) is shown as a bar plot with the length of each coloured bar corresponding to the time in that state. (B) Functional variation is important to understanding conscious processing. Mean functional activity (excitability) may vary from low to high. The integration and segregation of functional connectivity also informs the complexity of state information processing. We measure these using efficiency (global information sharing) and modularity (capacity for independent processing) which are interdependent, for instance the maximally integrated connectome is also minimally modular (top left), while other configurations exist somewhere in this space. (C) Spatiotemporal community analysis allows for states to be partitioned into functionally related modules that share members across temporally related states. These communities may be visualised by their fractional membership, the expected proportion of time a region spends in each community, here shown by colour opacity from light (low membership) to dark (high membership).

The information (or entropy) rate measures the expected information gained by sampling from a state space model over a sufficiently long state time Cover (1999). It provides an estimate of the temporal information capacity or complexity of the underlying process (see Supplementary Section S.8), differentiating it from the switching rate, which does not account for the diversity of probabilistically realisable states. As a result, a model can have high switching rate while having relatively low complexity (Fig. 2A). Brain state diversity was also measured directly using the entropy of the FO in order to consider subject-specific (rather than model or group-level) state diversity.

### Comparing States by Functional and Community Activity

HMM state functional connectivity and activity provide important modes to compare conditions of consciousness; brain activity may range from excitable (high absolute mean functional activity) to quiescent. We use paired distance correlation between states to compare them across both functional mean activity as well as functional connectivity, Fig. 2B. The states themselves encode complex connectivity relationships; these high dimensional relationships are often non-linear and in contrast to Pearson correlation, distance correlation takes into account such non-linear dependencies Geerligs, Henson, et al. (2016). Fig. 2B shows a range from dark blue (quiescent) to yellow (excitable) brain states. In addition, we apply the *signed distance correlation* from Pardo-Diaz et al. (2021) (with the sign being the sign of the standard Pearson correlation) to provide additional information on state dependencies.

High modularity and high efficiency are both necessary properties for the brain to perform complex tasks when integration of information from multiple sources or brain regions are involved Capouskova, Zamora-López, Kringelbach, and Deco (2023). The competing requirements for both appropriate segregation and integration of information processing need to be balanced in order to support both homeostasis and appropriate responsiveness to stimuli Deco, Tononi, Boly, and Kringelbach (2015); Jang, Mashour, Hudetz, and Huang (2024); R. Wang et al. (2021). Modularity is a proxy for segregation, with high modularity indicating a high capacity of the state to independently process multiple input streams, while high efficiency allows for fast simultaneous information sharing and integration between brain areas Meunier, Lambiotte, and Bullmore (2010); Sarasso et al. (2021); Zamora-López, Zhou, and Kurths (2010). The bottom of Fig. 2B shows how these two factors form a combined integration-segregation space within which functional reconfiguration occurs from state to state to meet ever changing informational processing demands on the brain.

The overall level of information integration within each state was assessed using the global efficiency of the state functional connectivity; the global efficiency measures how easily information can be propagated within a graph and is an established measure of functional integration in resting state fMRI Rubinov and Sporns (2010). Efficiency calculations were normalised against efficiency values for graphs with randomly permuted edge weights in order to control for overall graph connectivity and edge strength differences between states (see Supplementary Section S.9). Modules (indicated by node colours in Fig. 2B) here correspond to communities which were estimated using modularity maximisation Girvan and Newman (2002).

Fig. 2C sketches one of the primary outputs of our spatiotemporal community analysis. The fractional membership is the long run proportion of time that each brain region is expected to spend in a given community (see Supplementary Section S.10 for further details). To characterise these dynamic communities, we organise states and regions by performing hierarchical clustering using Ward’s algorithm Ward Jr (1963). This process is also used to uncover common spatiotemporal patterns of functional organisation across states and ROIs.

### Comparing Functional to Structural Connectivity

Patterns of functional organisation are dependent upon an underlying structural connectome Barttfeld et al. (2015); Demertzi et al. (2019); Luppi et al. (2023); Tan et al. (2019). In order to test whether deep anaesthesia or wakeful state functional connectivity is more related to structural connectivity, we compared the degree and level of integration of each state across conditions to the structural connectome using distance correlation. We also used the Adjusted Mutual Information (AMI) to quantify the similarity in community composition between structural and dynamic state models. The AMI is a measure of agreement between two partitions of brain regions that is more robust to differences in partition size than other measures such as the Adjusted Rand Index Romano, Vinh, Bailey, and Verspoor (2016) (see Supplementary Section S.11 and S.12). In order to closely compare structure and functional relationships at the level of individual brain regions, and in line with our we used the local efficiency first proposed in Rubinov and Sporns (2010). We employed the robust variant of the local efficiency proposed in Y. Wang, Ghumare, Vandenberghe, and Dupont (2017).

## RESULTS

### Selection of Models

We trained and analysed two dynamic models using data from each condition. The number of states for both wakeful and deep anaesthesia models was selected based on the maximum entropy of state occurrence (FO). This resulted in two different models that maximise the entropy, with 33 states for wakefulness, and 19 for anaesthesia, respectively (see Supplementary Fig. 2). We excluded sporadic states that featured in less than 20% of subject trajectories; this low threshold was chosen to allow for some subject-specific expression; see Supplementary Fig. 3 for the percentage of subjects visited in each state. The resulting models had *K_wake_*= 32 and *K_anaes_* = 18 states. For validation purposes, we also trained one combined model using both anaesthesia and wakeful data concatenated to produce a single model with the same number of initial states as wakefulness, *K_comb_*= 33, even after applying the same state exclusion criterion.

### Brain State Transition Graphs

Fig. 3 shows the transition probabilities between states as the edges in a directed graph of states (nodes) for both anaesthesia and wakefulness; wakefulness states are prefixed W-, while anaesthesia states are prefixed A-. In this figure, states are organised vertically by the level of brain state integration (as measured by normalised global efficiency) and the size of states is proportional to their sink centrality. The two graph models show clear separation along the (brain) state integration axis (*p ≤* 0.001 for the null hypothesis of global efficiency of the two models coming from the same distribution, Mann-Whitney U test, two-sided). The least integrated anaesthesia state is A-17; this is also the strongest sink state (see Supplementary Fig. 4 for the distribution of sink centralities relative to the strong sink threshold of 0.05), with A-17 being the most dominant at 0.38 (near to 40% of expected time in any state).

**Figure 3:**
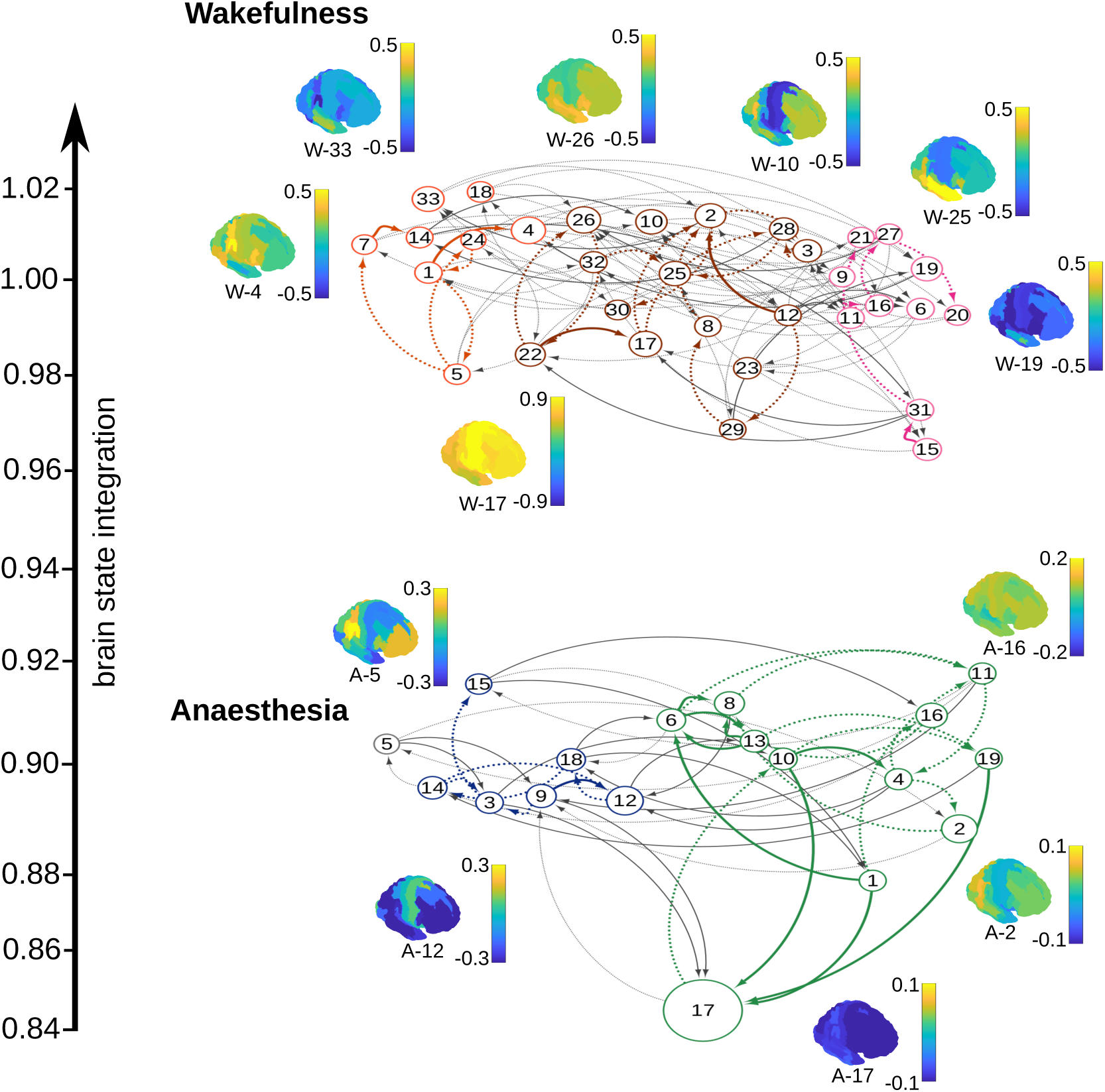
In the HMGM states are related by the probability of transition between them. This figure shows two graphs representing the respective state dynamics of subjects when awake or anaesthetised. Each node is a state the size of which is proportional to the state sink centrality. Edges represent that there is a non-negligible Transition Probability (TP) from one state to another. For visualisation purposes, only the two strongest edges from or to a node are shown with weaker probability edges (TP*<*0.05) denoted by dotted lines. The level of brain state integration (normalised global efficiency) is used to order the states on the y-axis, showing that anaesthesia states are significantly less integrated than wakeful states (*p ≤* 0.001, Mann-Whitney U test). States are organised horizontally into communities with coloured edges represent connections between states that share similar functional connectivity and dynamics. Here, we show brain maps for all states that account for 0.05 or more of the stationary distribution. We show brain maps of mean functional activity for all sink states (and associated colourbar), as well as state A-5 which is a transient outlier which does not belong to any cluster.

Fig. 3 shows that most sink states have many high probability in-edges. In particular, A-17 is reachable with relatively high probability from states within its own cluster, and is unlikely to be transitioned to from external states, except A-3 and A-9 which both act as bridges from the other cluster. Similarly, transitions to A-12, the second strongest sink state, generally occur from within its own cluster, but there are a few states in the A-17 dominated cluster that preferentially transition to A-12 acting as bridges between clusters. State A-5 has no strong connection to either cluster and has the lowest sink centrality of any state.

Wakeful dynamics are more diverse with no dominant sink state and three separate state clusters. Mean activity maps for sink states also show larger ranges (higher excitability) than in anaesthesia across all states (*p* = 0.017 *≤* 0.05 for the null hypothesis of activity being equal, measured by absolute mean activity, Mann-Whitney U test, two sided). Overall, the simpler modular structure of the anaesthesia model suggests lower complexity dynamics and indeed the anaesthesia information rate of 0.940 is less than the wakeful information rate of 1.82. Supplementary Fig. 5 shows that when varying the number of states in the model a difference between the two models persists, indicating that the difference in complexity is not simply due to a difference in the number of states but is due to the emergence of dominant states in anaesthesia that affect the reachability of other brain states.

### Sink State Dominance and Anaesthesia Quiescence

Fig. 4A and Fig. 4B show the stationary distribution across all subjects as pie charts with wedges representing sink states with sink centrality at least 0.05. Again, we see strong sink state dominance in anaesthesia and a more uniform stationary distribution in wakefulness. Fig. 4C and 4D show the top 50% most extreme regional activity for each sink state in wakefulness and anaesthesia, respectively. Nearly all regions shown for A-12 and A-17 have below-mean regional BOLD activity. In general, wakeful states have higher absolute mean activity across regions than in anaesthesia (*p ≤* 0.001, Mann-Whitney U test, two-sided). This also holds for the combined trial validation model (Supplementary Fig. 3) in which anaesthesia trials were dominated by a single state (state 29) with lower than baseline mean absolute functional activity.

**Figure 4:**
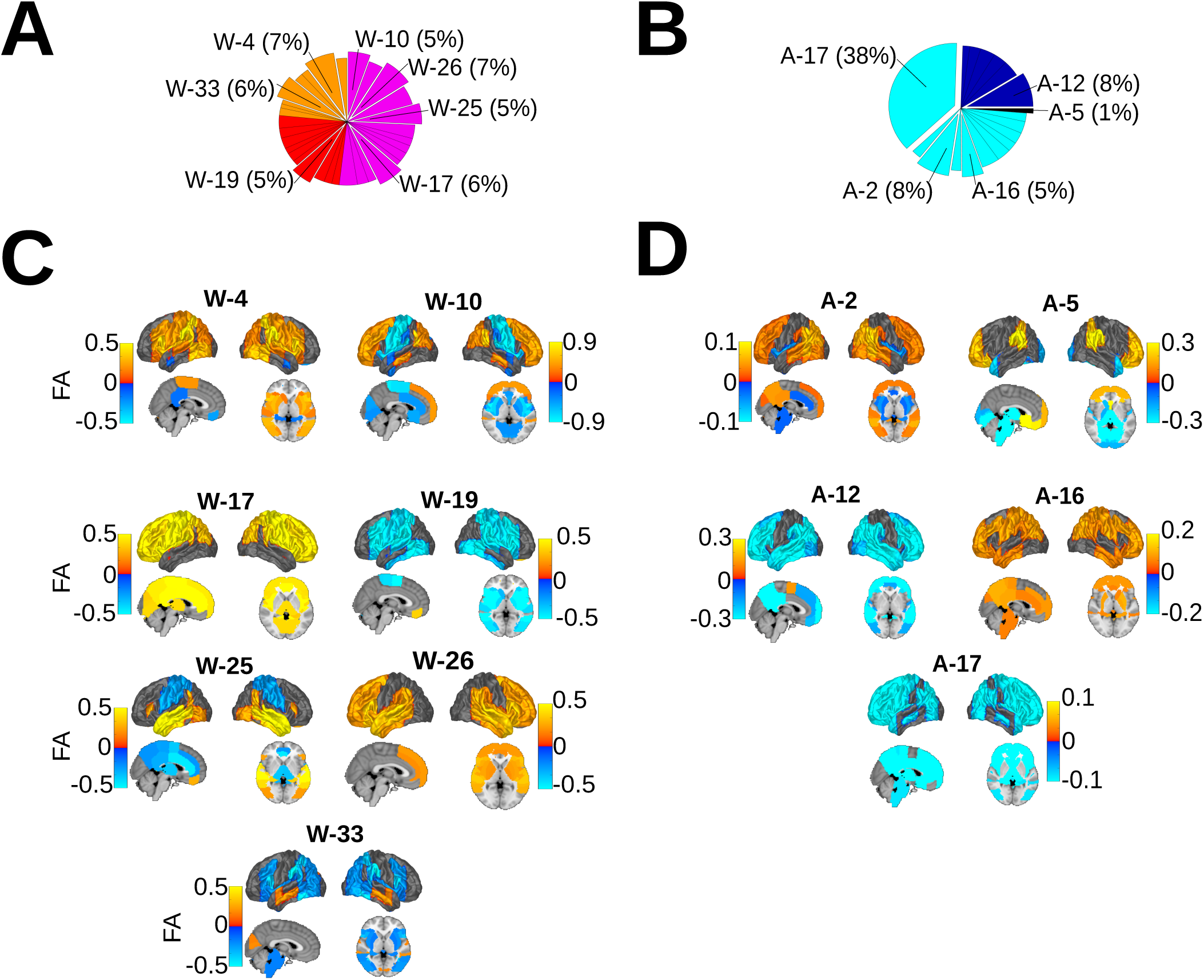
This figure summarizes the state activity of important states, highlighting the sink states as well as the outlying state A-5 and the prominent areas of surface Functional Activity (FA) for each. The stationary probability is shown for (A) wakefulness and (B) anaesthesia, with wedges indicating the sink states. In (C), the top 50% mean absolute functional activity in wakefulness is shown as a surface plot with a variety of high and low activity regions. (D) shows activity in the anaesthesia sink states, with much lower absolute mean functional activity across these states, indicative of lower absolute mean activity in anaesthesia as a whole (*p ≤* 0.001, Mann-Whitney U test, two-sided).

### Sink States Dominate Across Subjects in Anaesthesia but not Wakefulness

Fig. 5A shows the heterogeneity of state dynamics in wakefulness suggested in Fig. 3. While some states do seem more prevalent amongst certain subject groups, e.g. sinks states W-4 and W-26 can be differentiated from the subject-specific patterns of sink states W-2 and W-33, no single state dominates any one grouping. By contrast, Fig. 5B shows that subjects under deep anaesthesia can be separated into two groups depending on sink state dominance (see Supplementary Fig. 4 for relative sink centralities across states and conditions). The majority of subjects are dominated by A-17, and the remainder have a relatively high A-12 occupancy or are occupied by subject-specific sink states (A-2 and A-16). There does not appear to be significant overlap between anaesthesia and wakefulness in how subjects are grouped. It is unclear why not all subjects share the same dominant state; however, we note that the latter group is significantly younger (*p* = 0.026 *<* 0.05, Mann-Whitney U test). No significant difference in age was found between the two groups separating wakeful subjects by state FO (*p* = 0.264 *>* 0.05, Mann-Whitney U test).

**Figure 5:**
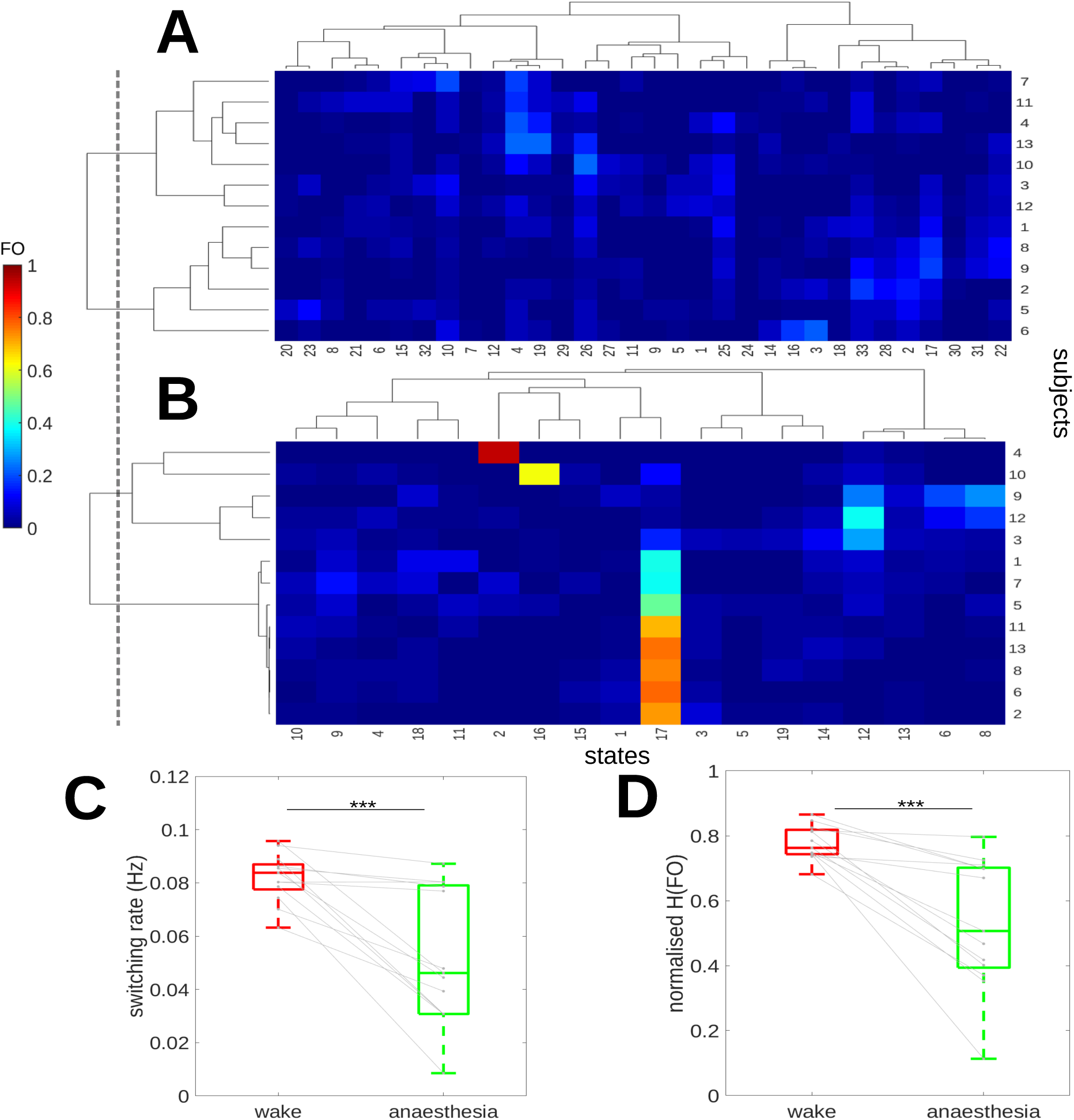
Subject specific state dynamics is shown for both anaesthesia and wakefulness. The dotted line separates both subjects into clusters according to FO correlation. (A) shows the FO heat map for wakefulness as a clustergram where subjects (rows) and states (columns) have been hierarchically clustered by Pearson correlation using Ward’s algorithm. (B) In anaesthesia, dynamics tend to be dominated by a subset of sink states, including subjects 4 and 10 which show a strong preference for A-2 and A-16 respectively. (C) Shows a boxplot comparing wakeful and and anaesthetised switching rates and normslised occupancy entropy (H(FO)). The state switching rate (Hz) is more varied but significantly lower in anaesthesia (*p ≤* 0.001, Wilcoxon Signed-rank test, two-sided). The H(FO), displayed in (D), shows a similar pattern to the switching rate with highly variable but overall lower state diversity present in anaesthetised versus awake subjects (*p ≤* 0.001, Wilcoxon Signed-rank test).

The FO of the two strongest states (A-12 and A-17) have a strong negative Pearson correlation of *ρ* = *−*0.922 (*p ≤* 0.01, permutation test; see Supplementary Section S.13 for details). Together A-12 and A-17 account for *≥* 50% of the mean FO across all subjects in deep anaesthesia, indicating that a majority of subject time is spent in one of these two states, with the overwhelming majority of time spent in A-17. This is in comparison to wakefulness in which no two states can account for more than 20% of mean FO. In the validation models using either the combined data (including both wakeful and anaesthesia trials in a single model) or an alternative model with *K* = 20 states, anaesthesia trials were also dominated by sink states (states 29 and 20 respectively, see Supplementary Fig. 6 and 7).

Fig. 5C shows state switching rates across subjects in both conditions, with higher switching observed in wakefulness (*p ≤* 0.001, Wilcoxon Signed-rank test, two sided). Fig. 5D shows state diversity, as measured by the normalised entropy of the FO, with higher state diversity in wakeful subjects when compared to anaesthesia (*p ≤* 0.001, Wilcoxon Signed-rank test, two sided). Together these results suggest that dynamics under deep anaesthesia are not only simpler but also less dynamic, with a lower fraction of states participating in the dynamics of most subjects, as would be expected for dynamics with a lower information rate. However, there is also some evidence that subjects are more separable, for instance by state dominance, likely due to the subject-specific anaesthesia depth, which varies even when subjects are held at the same ESC, and which may depend on subject-specific age and susceptibility factors Nunes, Alonso, Castro, Amorim, and Mendonça (2007).

### Stratification of Functional Connectivity in Anaesthesia

In the wakeful condition, state diversity results in highly diverse spatiotemporal community reorganisation. Hierarchical clustering of spatiotemporal community membership, combining functional and temporal relationships, reveal highly distributed dynamics, with each community including diverse and dynamically changing membership (see Fig. 6A). These communities dynamically represent shifts between resting state networks including visual, salience-default mode and sensory networks respectively. The dynamics is visible in Fig. 6B where regional switching between communities is reflected in the mosaic pattern of community associations.

**Figure 6:**
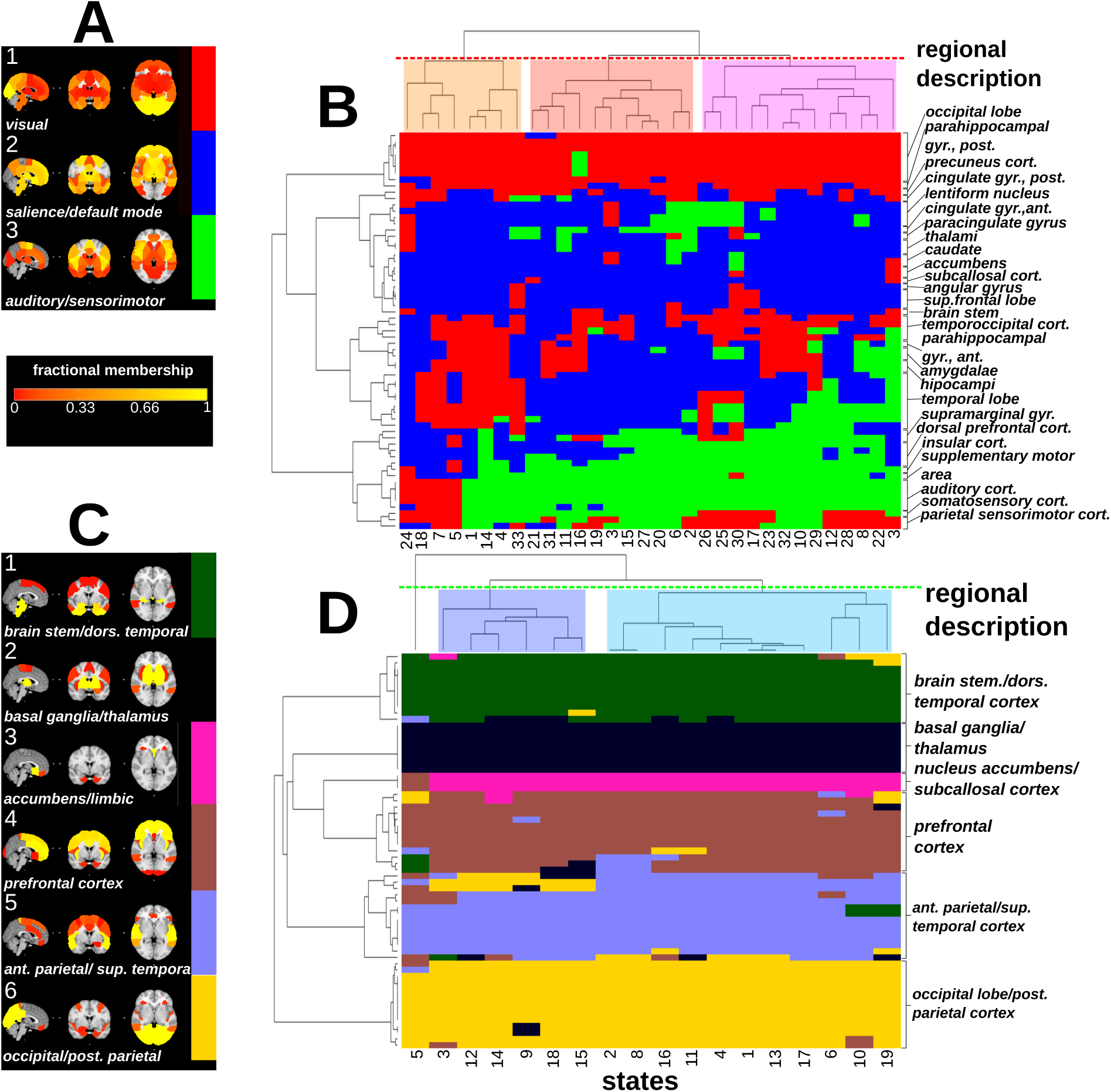
This figure summarises the results of spatiotemporal consensus community detection across regions (rows) and states (columns). Each ROI is coloured according to its community membership. ROIs and states have been hierarchically clustered in both deep anaesthesia and wakefulness using the Jaccard index similarity so that states with similar community composition and regions with similar membership are co-clustered (single colour patches). (A) this plot shows the regional composition of each wakeful community as a brain map where colour represent the fractional membership of the region. Communities have been broadly described according to the text labels. (B) the clustergram of wakeful communities shows a heterogeneous pattern of changing community membership. (C) shows brain maps for the deep anaesthesia condition. Brain regions are largely static (fractional membership near to one) and have a relatively restricted anatomically organisation with more total communities. (D) the anaesthesia clustergram shows regional stratification with minimal reorganisation.

The anaesthesia model (see Fig. 6C and D) reveals strikingly different results, with a greater number of less dynamic communities that are primarily restricted to closely anatomically related ROIs. There is far less change of membership between states in anaesthesia in comparison to wakefulness, resulting in stratified and near static dynamic reorganisation. Even though we see less switching in anaesthesia dynamics, sporadic co-recruitment of regions commonly associated with the so-called Resting State Networks does occur, for example in A-16 (exhibiting Default Mode-like activity in community 3).

Furthermore, the spatiotemporal hierachical clustering of communities in anaesthesia suggests that subcortical areas, most notably the thalamus and neighbouring regions (including the caudate), generally thought of as an information bottleneck for sensory inputs reaching the brain Alitto and Usrey (2003); Luppi et al. (2024), are isolated from cortical brain regions in almost all anaesthesia states. This is not seen in wakefulness, in which thalamic regions share and switch membership with multiple cortical regions depending on state, supporting the dynamic role of the thalamus in conscious homeostasis.

### Sink State Similarity Across Conditions

Plots of the functional connectome of each of the sink states are shown in Fig. 7A (thresholded to the top 5% strongest connections by magnitude for illustrative purposes), with mean activity maps for each state shown vertically. The heatmaps represent the pairwise distance correlation (displaying only strong relationships, *>* 0.5) between states across both conditions. Very few wakeful states share a significant relationship with any anaesthesia state on the basis of either functional activity (bottom left) or connectivity (top right). Fig. 7B shows that pairwise distance correlation of functional connectivity between wakeful states is significantly higher than in anaesthesia (*p ≤* 0.001, Mann-Whitney U test, two-sided). This difference in state self-similarity holds even when considering distance correlation in signed functional (Pearson) correlation between ROIs (*p ≤* 0.001 Mann-Whitney U test, two-sided). There is evidence for the suggestion that most anaesthesia states resemble neither each other nor are directly related to wakeful states, even when considering possible changes in the sign of functional relationships. A similar but weaker finding holds for functional activity (*p ≤* 0.001 Mann-Whitney U test, two-sided) with mean functional activity distance correlation lower in anaesthesia than in wakefulness, see Fig. 7C. Although anaesthesia states are generally superficially dissimilar, signed distance correlation does suggest that A-17 is at least somewhat positively related to all other anaesthesia states; this relationship between anaesthesia states makes it plausible to observe the perhaps surprising similarity in spatiotemporal community composition seen across anaesthesia states.

**Figure 7:**
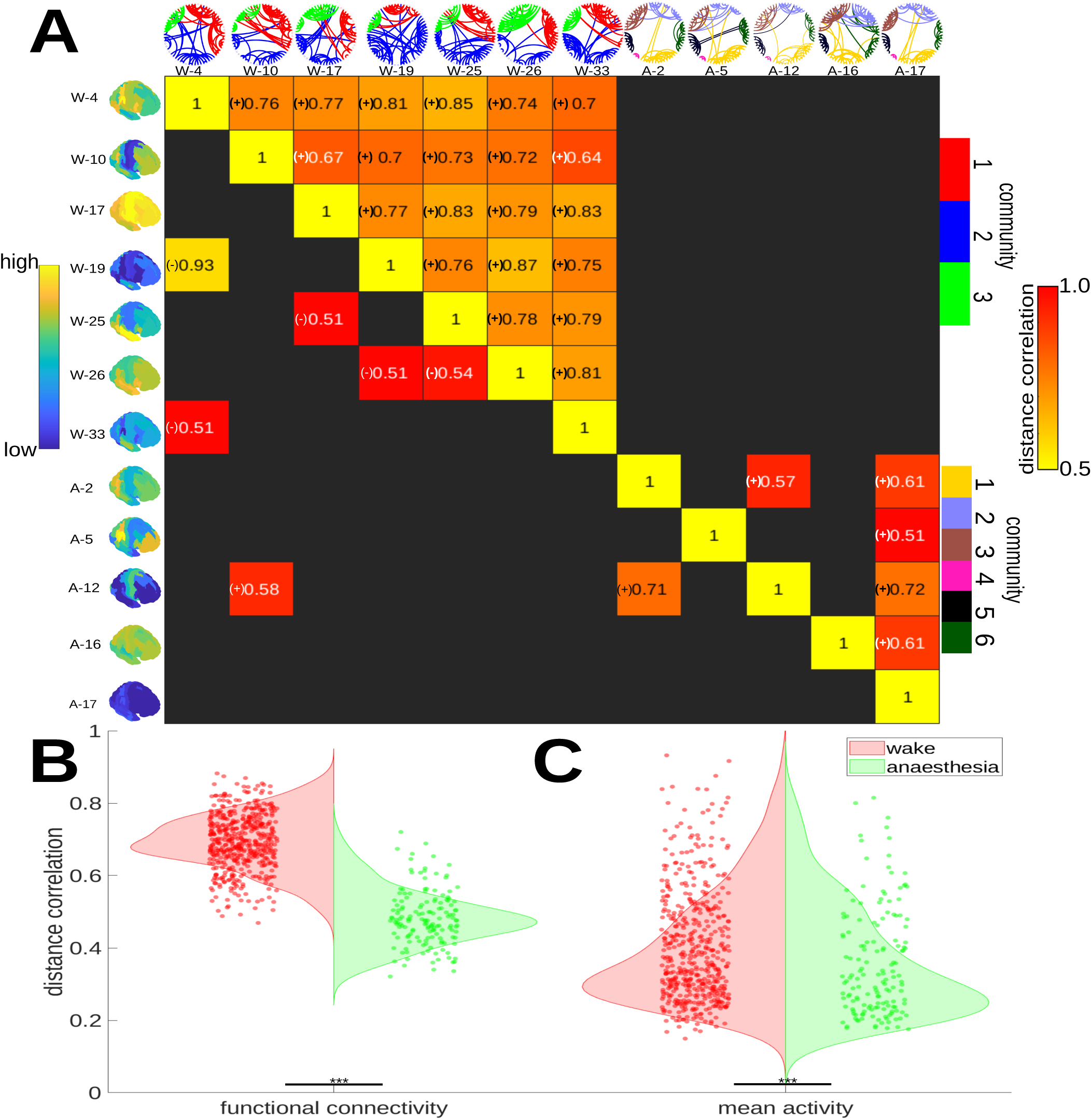
This figure illustrates the functional mean activity and connectivity across sink states in anaesthesia and wakefulness. (A) This figure is a heatmap of the pairwise signed distance correlation between sink states functional activity (bottom left) and sink state connectivity (top right). Brain maps (left) are the same as those in Fig. 3. Community interconnectivity (top) is shown as a circular graph, depicting the level of intercommunity connectivity in each state. (B) This violinplot shows the distance correlation in functional connectivity between pairs of states (dots) in (red) wakefulness and (green) anaesthesia (Mann-Whitney U, *p ≤* 0.001, two sided). Crosses denote outlying states. (C) Signed distance correlation is higher for mean activity between wakeful states though the difference appears smaller (Mann-Whitney U, *p ≤* 0.001, two-sided).

### Structure-Function Relationships in Wakefulness and Anaesthesia

Similarity analysis of the functional connectivity of each state with the underlying structural connectivity reveals significant differences in how wakeful and anaesthesia states relate to or agree with (under a given similarity measure) structural connectivity. The connectivity violinplot in Fig. 8 shows the distance correlation (dcor) between edge weights of a state and structural edge weights (each expressed as vectors). The community violinplot in Fig. 8 shows the results of comparing structural and functional modules in both conditions, as determined by consensus multi-resolution community analysis (Supplementary Fig. 8 for these modules). Here, we used the AMI, a standard measure of community agreement, to determine similarity in community organisation between brain states and structural organisation. Anaesthesia community composition and connectivity are moderately consistent with the observed structural modularisation, having an average AMI of 0.51 between states, when compared to wakefulness, having an AMI of 0.24 (*p ≤* 0.001, permutation test); we also validated these results using the ARI (see Supplementary Section S.11 and S.12). Wakefulness was found to be more correlated to structure in terms of local efficiency (integration) and degree centrality statistics (*p ≤* 0.001, Mann-Whitney U, two-sided). The connectivity and degree centrality results hold even when signed Pearson correlation is used in place of the unsigned (absolute) connectivity we use in our framework (*p ≤* 0.001, Mann-Whitney U, two-sided).

**Figure 8:**
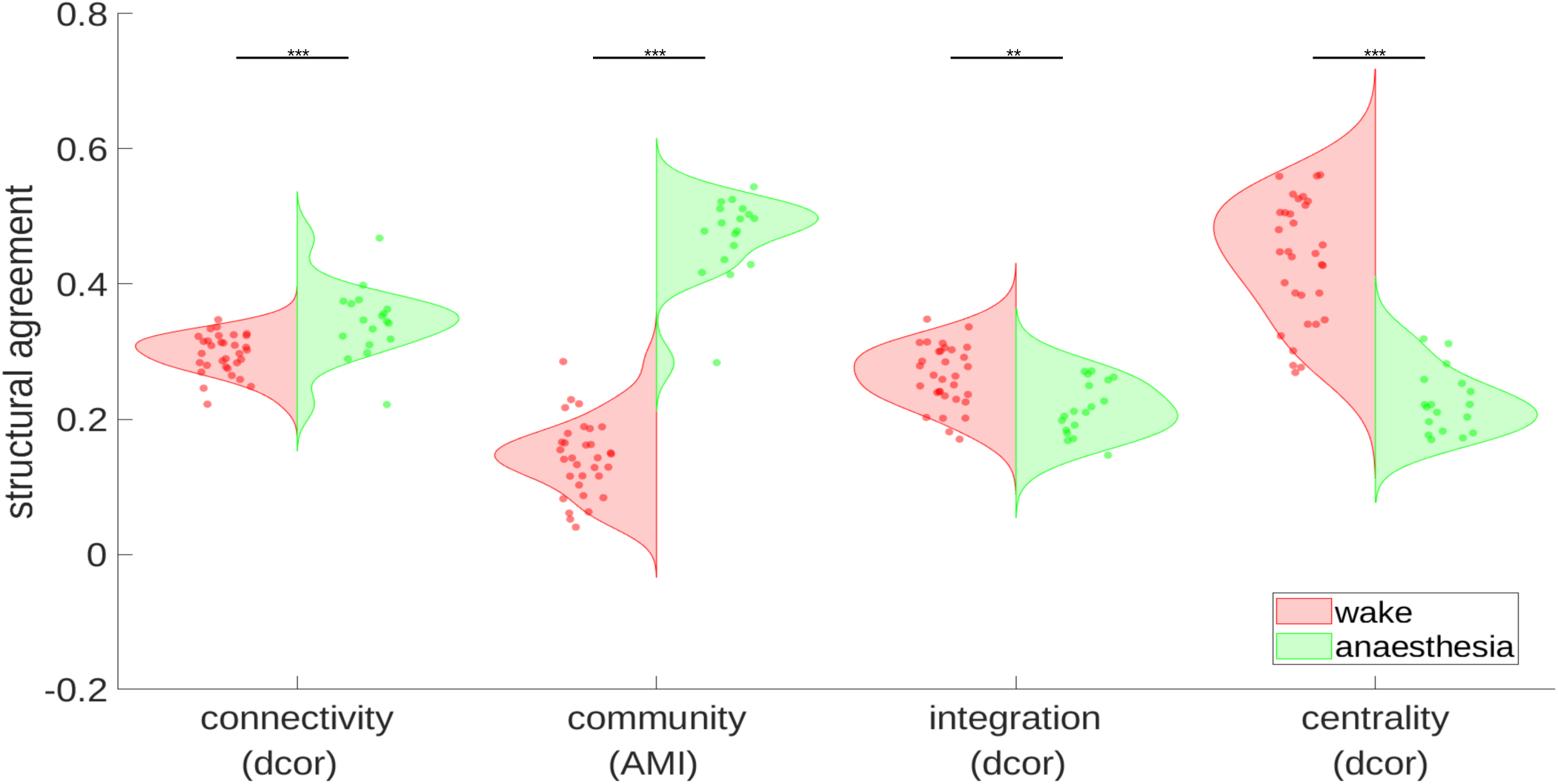
This figure shows the differences between brain state functional connectivity in wakefulness and anaesthesia as they relate to structural connectivity using either distance correlation (dcor) or AMI, visualised as a violinplot with states represented as either red (wakefulness) or green (anaesthesia) dots. The connectivity plot shows the relationship between brain state edge weights and structural network edge weights, in (red) wakefulness and (green) anaesthesia with anaesthesia state connectivity being more similar to structural connectivity (*p* = 0.001, Mann-Whitney U test, two-sided). The community violinplot shows the AMI, a measure of modular similarity, between brain state modules and structural modules suggesting that anaesthesia functional modules are more similar to the normative structural modularisation of the brain than more dynamic wakeful states (*p <* 0.001, permutation test). For local comparison, integration is here measured by the local efficiency compared to the structural local efficiency (*p ≤* 0.001, Mann-Whitney U test, two-sided). Centrality shows the distance correlation in degree centrality between structure and state graphs (*p ≤* 0.001, Mann-Whitney U test, two-sided).

### Sink State Occupancy Correlates with Age and Susceptibility

Given the overwhelmingly dominant role of the strongest sink state in determining the range of anaesthesia brain state dynamics, it is important to determine which factors, if any, are most determinative of sink state dominance. Multivariate linear regression was performed to determine which factors most closely predict the time subjects spent (FO) in the most dominant sink state A-17. Increasing age was found to be the single variable most associated with positive increases in A-17 FO (*p* = 0.005 *≤* 0.01, t-test, two-sided) when considered alone. However, time at loss of responsiveness (*T_LOBR_*) in combination with age was found to be the best performing predictor of A-17 FO with respect to both the BIC (see Supplementary Table 2) and adjusted R^2^ criteria, with significant improvements in goodness of fit over the age-only model (*p* = 0.040 *≤* 0.05, Chi-squared test). The model shows that age is a significant predictor (*p* = 0.004 *≤* 0.01, t-test, two-sided), while *T_LOBR_* is marginally negatively associated with A-17 FO (*p* = 0.079, t-test, two-sided). Supplementary Fig. 9 shows the alternative model which includes time conditioned on age. Surprisingly, grey matter density (see Supplementary Section S.14) did not add any significant model improvement (see Supplementary Tables 3 to 6).

## DISCUSSION

In this paper we contrast the dynamics of resting wakefulness and unconsciousness under deep anaesthesia using data from a multi-phase study of subjects undergoing propofol-induced anaesthesia, focusing on analysis of fMRI BOLD activity. In the analysis we apply a combined HMM-based dynamic state space model Vidaurre et al. (2016), as well as a new spatiotemporal community detection approach, in order to determine the dynamic modular reorganisation of brain activity. We provide results corroborating a general decreases in temporal complexity during unconsciousness seen in other investigations Barttfeld et al. (2015); Demertzi et al. (2019); Mediano et al. (2023), and propose that this dynamic may be due to the dominance of low complexity sink states and widespread functional disintegration. We divide these results into three primary findings which we discuss here.

First, regarding the brain state dynamics, we corroborate and expand on existing research showing that unconscious dynamics Mediano et al. (2023); Stevner et al. (2019) and anaesthesia dynamics in particular Barttfeld et al. (2015); Demertzi et al. (2019); Hudetz, Liu, and Pillay (2015); Liang et al. (2024), are temporally less complex both in terms of the effective diversity and reachability of states and in state switching. We suggest that the attraction of anaesthesia dynamics towards a small set of sink states, with a severely reduced capacity for integration and cortical and subcortical excitability, may, in part, be responsible for this reduction in complexity. However, visits to other states may suggest moments of brief burst suppression, partial arousal in responses to external or internal perturbations Dueck et al. (2005); Mhuircheartaigh et al. (2013); Z. Zhang et al. (2019), or even commonly reported dream-like events Hack et al. (2024). Sink state dominance seems much less prevalent in wakefulness with comparatively much weaker attractor-like states, perhaps owing to a need for higher levels of responsiveness.

Second, we observe fragmentation of functional modules under anaesthesia into small, static subnetworks with little thalamocortical or even cortical-subcortical re-organisation from state to state. In particular, thalamic community associations mostly occurred between neighbouring subcortical and entorhinal regions. Notably, some interchange is observed between prefrontal, parietal and nucleus accumbens, suggesting some sporadic default mode-like activity. This thalamocortical isolation has also been shown from static measures of functional connectivity Mhuircheartaigh et al. (2013). Deteriorated but persistent default mode network activity has been observed in other studies of anaesthesia functional connectivity, with the anterior portions being most affected Luppi et al. (2019). In our model, anaesthesia is contrasted with wakefulness in which larger communities dynamically exchange members between states to facilitate more complex task performance, required for conscious awareness and for responsiveness to the external and internal environment Barttfeld et al. (2015). This result may be surprising as we find that anaesthesia state connectivity was generally less self-similar than in wakefulness. However, this is consistent with the hypothesis that there is a persistent global core to functional connectivity in wakefulness Onoda and Akama (2023), likely including the thalamus as an important hub Luppi et al. (2024); Mhuircheartaigh et al. (2013); Tasserie et al. (2022). Stratification may result in this core being partially fragmented into anatomically restricted regions during unconsciousness Hu, Wang, Zhang, Luo, and Wang (2024).

Third, we examine the dynamic structure-function relationship in relation to both connectvity and modularistion. Our results support the generally observed increase in similarity between functional and structural connectivity as measured by connection weights Barttfeld et al. (2015); Demertzi et al. (2019), as well as providing new evidence of weak similarity between dynamic modularisation and structural modularisation under anaesthesia. In particular, we note that most anaesthesia communities appear spatially constrained to specific brain areas, perhaps reflecting propagation of anaesthesia signal along short-range structural connections that then fail to propagate more widely as observed in Gosselin, Bagur, and Bathellier (2024); Mhuircheartaigh et al. (2013). Long-range structural connections have been shown to be sparser and more targeted in-comparison to short-range connections Betzel and Bassett (2018), supporting diverse, complex functional processes in wakefulness that may be absent in unconsciousness and in severe cases of DOCs Altmayer et al. (2024); Luppi et al. (2021, 2025) In addition to anatomical modularisation, this difference between short and long-range connectivity may explain the dissimilarity in efficiency and degree centrality between anaesthesia states and underlying structural connectivity.

Finally we indicate some implications of our study regarding patient-specific targeting of anaesthesia depth. The strong attractor-like dynamics towards low complexity, low excitability anaesthesia states which we observe reflect the imperturbability of subjects from anaesthesia. The association of specific brain states with levels of arousal under anaesthesia has previously been reported Barttfeld et al. (2015). We demonstrate subject-specific sink state activity with changes in the degree of sink state dominance and the specific sink state active, defining possible differences in the steady state for anaesthesia homeostasis which may indicate differing capacities for information integration and arousal. We find a tentative link between sink state dominance and anaesthetic susceptibility, specifically the age of the subject, and, to a lesser extent, time at loss of responsiveness. This may relate to previous evidence of age-related state activity in surgical EEG Li et al. (2020). Furthermore, these results suggest that sink state dominance may share similar features with common monitors of anaesthesia depth that are also known to be modulated by age and susceptibility-specific factors Cortínez et al. (2008); Obert et al. (2021); Warnaby, Sleigh, Hight, Jbabdi, and Tracey (2017). More tentatively, the persistence of sink state-like dynamics may be a possible future marker for recovery from DOCs.

With regards to limitations, our study was limited in its structure-function comparisons by the availability of patient-specific structural data that would enable us to make subject-specific comparisons. Finding a parcellation that was computationally tractable and relevant in a functional context was prioritised partially due to the relatively small cohort size for the ultra-slow anaesthethesia study. Due to this choice, some cortical areas lacked comparable left and right regions, preventing us from observing possible lateralisation of activity in some cases. Sample size and small age range may also explain the apparent weak association between grey matter volume and age Terribilli et al. (2011), and limit the generalisability of the result to older ages who see the most significant detrimental affects of over anaesthesia Sevenius Nilsen et al. (2019); Wu et al. (2019). In this regard, further experimental studies would be of interest.

The focus on brain state has allowed for the development of new ways of analysing the complexity of the brain in action Barnett and Seth (2023); Barttfeld et al. (2015); Mediano et al. (2022). We believe that new complexity measures should put special focus on combining both spatial and temporal complexity, while taking into account the brain’s dynamic capacity for integrating the interdependent streams of information that support complex behaviour. Our HMGM approach provides a novel graphical solution to address the problem by decomposing the entire neural system into spatiotemporal subnetworks that capture the reorganisation necessary to sustain complex activity.

## ACKNOWLEDGMENTS

The authors are particularly grateful to Marco Fabus for his assistance in identifying periods of burst suppression from subject EEG. Minh Tran for his helpful advice in model design and testing. The authors are grateful to the following funding bodies: GR is funded in part by the UKRI EPSRC grant EP/T018445/1, EP/R018472/1, EP/X002195/1 and EP/Y028872/1, JW is funded by the UKRI BBSRC grant BB/Y003020/1 and KW acknowledges funding from the UKRI MRC grant MR/R006423/1. For the purpose of Open Access, the authors note that a CC BY public copyright licence applies to any Author Accepted Manuscript version arising from this submission.

## SUPPORTING INFORMATION

Supporting information including supporting figures and method details are included in the Supplementary Information document.

## REFERENCES

Aceto, P., Perilli, V., Lai, C., Sacco, T., Ancona, P., Gasperin, E., & Sollazzi, L. (2013). Update on post-traumatic stress syndrome after anesthesia. European Review for Medical & Pharmacological Sciences, 17(13).

Alitto, H. J., & Usrey, W. M. (2003). Corticothalamic feedback and sensory processing. Current Opinion in Neurobiology, 13(4), 440–445.

Altmayer, V., Sangare, A., Calligaris, C., Puybasset, L., Perlbarg, V., Naccache, L., … Rohaut, B. (2024). Functional and structural brain connectivity in disorders of consciousness. Brain Structure and Function, 1–14.

Bai, Y., He, J., Xia, X., Wang, Y., Yang, Y., Di, H., … Ziemann, U. (2021). Spontaneous transient brain states in EEG source space in disorders of consciousness. NeuroImage, 240, 118407.

Barnett, L., & Seth, A. K. (2023). Dynamical independence: discovering emergent macroscopic processes in complex dynamical systems. Physical Review E, 108(1), 014304.

Barttfeld, P., Uhrig, L., Sitt, J. D., Sigman, M., Jarraya, B., & Dehaene, S. (2015). Signature of consciousness in the dynamics of resting-state brain activity. Proceedings of the National Academy of Sciences, 112(3), 887–892.

Betzel, R. F., & Bassett, D. S. (2018). Specificity and robustness of long-distance connections in weighted, interareal connectomes. Proceedings of the National Academy of Sciences, 115(21), E4880–E4889.

Bonhomme, V., Staquet, C., Montupil, J., Defresne, A., Kirsch, M., Martial, C., … others (2019). General anesthesia: a probe to explore consciousness. Frontiers in Systems Neuroscience, 13, 36.

Brin, S., & Page, L. (1998). The anatomy of a large-scale hypertextual web search engine. Computer Networks and ISDN Systems, 30(1-7), 107–117.

Bryson, G. L., & Wyand, A. (2006). Evidence-based clinical update: general anesthesia and the risk of delirium and postoperative cognitive dysfunction. Canadian Journal of Anesthesia, 53(7), 669.

Capouskova, K., Zamora-López, G., Kringelbach, M. L., & Deco, G. (2023). Integration and segregation manifolds in the brain ensure cognitive flexibility during tasks and rest. Human brain mapping, 44(18), 6349–6363.

Castro, P., Luppi, A., Tagliazucchi, E., Perl, Y. S., Naci, L., Owen, A. M., … Cofré, R. (2024). Dynamical structure-function correlations provide robust and generalizable signatures of consciousness in humans. Communications Biology, 7(1), 1224.

Cortínez, L. I., Trocóniz, I. F., Fuentes, R., Gambus, P., Hsu, Y.-W., Altermatt, F., & Munoz, H. R. (2008). The influence of age on the dynamic relationship between end-tidal sevoflurane concentrations and bispectral index. Anesthesia & Analgesia, 107(5), 1566–1572.

Cover, T. M. (1999). Elements of information theory. John Wiley & Sons.

Deco, G., Tononi, G., Boly, M., & Kringelbach, M. L. (2015). Rethinking segregation and integration: contributions of whole-brain modelling. Nature reviews neuroscience, 16(7), 430–439.

Demertzi, A., Tagliazucchi, E., Dehaene, S., Deco, G., Barttfeld, P., Raimondo, F., … others (2019). Human consciousness is supported by dynamic complex patterns of brain signal coordination. Science Advances, 5(2), eaat7603.

Desikan, R. S., Ségonne, F., Fischl, B., Quinn, B. T., Dickerson, B. C., Blacker, D., … others (2006). An automated labeling system for subdividing the human cerebral cortex on mri scans into gyral based regions of interest. Neuroimage, 31(3), 968–980.

Dueck, M., Petzke, F., Gerbershagen, H., Paul, M., Hesselmann, V., Girnus, R., … others (2005). Propofol attenuates responses of the auditory cortex to acoustic stimulation in a dose-dependent manner: A fmri study. Acta anaesthesiologica scandinavica, 49(6), 784–791.

Fan, S.-Z., Yeh, J.-R., Chen, B.-C., Shieh, J.-S., et al. (2011). Comparison of EEG approximate entropy and complexity measures of depth of anaesthesia during inhalational general anaesthesia. Journal of Medical and Biological Engineering, 31(5), 359–366.

Forrest, F., Tooley, M., Saunders, P., & Prys-Roberts, C. (1994). Propofol infusion and the suppression of consciousness: the EEG and dose requirements. BJA: British Journal of Anaesthesia, 72(1), 35–41.

Fritz, B. A., Kalarickal, P. L., Maybrier, H. R., Muench, M. R., Dearth, D., Chen, Y., … Avidan, M. S. (2016). Intraoperative electroencephalogram suppression predicts postoperative delirium. Anesthesia and Analgesia, 122(1), 234.

Galatolo, S., Hoyrup, M., & Rojas, C. (2010). Effective symbolic dynamics, random points, statistical behavior, complexity and entropy. Information and Computation, 208(1), 23–41.

Geerligs, L., Henson, R. N., et al. (2016). Functional connectivity and structural covariance between regions of interest can be measured more accurately using multivariate distance correlation. NeuroImage, 135, 16–31.

Girvan, M., & Newman, M. E. (2002). Community structure in social and biological networks. Proceedings of the national academy of sciences, 99(12), 7821–7826.

Gosselin, E., Bagur, S., & Bathellier, B. (2024). Massive perturbation of sound representations by anesthesia in the auditory brainstem. Science Advances, 10(42), eado2291.

Hack, L. M., Sikka, P., Zhou, K., Kawai, M., Chow, H. S., & Heifets, B. (2024). Reduction in trauma-related symptoms after anesthetic-induced intra-operative dreaming. American Journal of Psychiatry, 181(6), 563–564.

He, M., Das, P., Hotan, G., & Purdon, P. L. (2023). Switching state-space modeling of neural signal dynamics. PLoS Computational Biology, 19(8), e1011395.

Heine, L., Soddu, A., Gómez, F., Vanhaudenhuyse, A., Tshibanda, L., Thonnard, M., … Demertzi, A. (2012). Resting state networks and consciousness: alterations of multiple resting state network connectivity in physiological, pharmacological, and pathological consciousness states. Frontiers in Psychology, 3, 295.

Horn, A., & Blankenburg, F. (2016). Toward a standardized structural–functional group connectome in mni space. NeuroImage, 124, 310–322.

Hu, Y., Wang, Y., Zhang, L., Luo, M., & Wang, Y. (2024). Neural network mechanisms underlying general anesthesia: Cortical and subcortical nuclei. Neuroscience Bulletin, 1–17.

Hudetz, A. G., Liu, X., & Pillay, S. (2015). Dynamic repertoire of intrinsic brain states is reduced in propofol-induced unconsciousness. Brain Connectivity, 5(1), 10–22.

Ito, T., Yang, G. R., Laurent, P., Schultz, D. H., & Cole, M. W. (2022). Constructing neural network models from brain data reveals representational transformations linked to adaptive behavior. Nature Communications, 13(1), 673.

Jang, H., Mashour, G. A., Hudetz, A. G., & Huang, Z. (2024). Measuring the dynamic balance of integration and segregation underlying consciousness, anesthesia, and sleep in humans. Nature communications, 15(1), 9164.

Jenkinson, M., Beckmann, C. F., Behrens, T. E., Woolrich, M. W., & Smith, S. M. (2012). Fmrib software library (FSL). NeuroImage, 62(2), 782–790.

Lancichinetti, A., & Fortunato, S. (2012). Consensus clustering in complex networks. Scientific Reports, 2(1), 1–7.

Leroux, B. G. (1992). Maximum-likelihood estimation for hidden markov models. Stochastic processes and their applications, 40(1), 127–143.

Li, D., & Mashour, G. A. (2019). Cortical dynamics during psychedelic and anesthetized states induced by ketamine. Neuroimage, 196, 32–40.

Li, D., Puglia, M. P., Lapointe, A. P., Ip, K. I., Zierau, M., McKinney, A., & Vlisides, P. E. (2020). Age-related changes in cortical connectivity during surgical anesthesia. Frontiers in Aging Neuroscience, 11, 371.

Li, D., Vlisides, P. E., & Mashour, G. A. (2022). Dynamic reconfiguration of frequency-specific cortical coactivation patterns during psychedelic and anesthetized states induced by ketamine. Neuroimage, 249, 118891.

Liang, Z., Tang, B., Chang, Y., Wang, J., Li, D., Li, X., & Wei, C. (2024). State-related electroencephalography microstate complexity during propofol-and esketamine-induced unconsciousness. Anesthesiology, 140(5), 935–949.

Liu, Z., Si, L., Xu, W., Zhang, K., Wang, Q., Chen, B., & Wang, G. (2022). Characteristics of eeg microstate sequences during propofol-induced alterations of brain consciousness states. IEEE Transactions on Neural Systems and Rehabilitation Engineering, 30, 1631–1641.

López-González, A., Panda, R., Ponce-Alvarez, A., Zamora-López, G., Escrichs, A., Martial, C., … others (2021). Loss of consciousness reduces the stability of brain hubs and the heterogeneity of brain dynamics. Communications Biology, 4(1), 1037.

Luppi, A. I., Craig, M. M., Coppola, P., Peattie, A. R., Finoia, P., Williams, G. B., … Stamatakis, E. A. (2021). Preserved fractal character of structural brain networks is associated with covert consciousness after severe brain injury. NeuroImage: Clinical, 30, 102682.

Luppi, A. I., Craig, M. M., Pappas, I., Finoia, P., Williams, G. B., Allanson, J., … others (2019). Consciousness-specific dynamic interactions of brain integration and functional diversity. Nature communications, 10(1), 4616.

Luppi, A. I., Golkowski, D., Ranft, A., Ilg, R., Jordan, D., Bzdok, D., … others (2025). General anaesthesia decreases the uniqueness of brain functional connectivity across individuals and species. Nature Human Behaviour, 1–18.

Luppi, A. I., Uhrig, L., Tasserie, J., Signorelli, C. M., Stamatakis, E. A., Destexhe, A., … Cofre, R. (2024). Local orchestration of distributed functional patterns supporting loss and restoration of consciousness in the primate brain. Nature communications, 15(1), 2171.

Luppi, A. I., Vohryzek, J., Kringelbach, M. L., Mediano, P. A., Craig, M. M., Adapa, R., … others (2023). Distributed harmonic patterns of structure-function dependence orchestrate human consciousness. Communications biology, 6(1), 117.

Marshall, W., Gomez-Ramirez, J., & Tononi, G. (2016). Integrated information and state differentiation. Frontiers in Psychology, 7, 926.

Mediano, P. A., Rosas, F. E., Luppi, A. I., Jensen, H. J., Seth, A. K., Barrett, A. B., … Bor, D. (2022). Greater than the parts: a review of the information decomposition approach to causal emergence. Philosophical Transactions of the Royal Society A, 380(2227), 20210246.

Mediano, P. A., Rosas, F. E., Luppi, A. I., Noreika, V., Seth, A. K., Carhart-Harris, R. L., … Bor, D. (2023). Spectrally and temporally resolved estimation of neural signal diversity. bioRxiv, 2023–03.

Meunier, D., Lambiotte, R., & Bullmore, E. T. (2010). Modular and hierarchically modular organization of brain networks. Frontiers in Neuroscience, 4, 200.

Mhuircheartaigh, R. N., Warnaby, C., Rogers, R., Jbabdi, S., & Tracey, I. (2013). Slow-wave activity saturation and thalamocortical isolation during propofol anesthesia in humans. Science Translational Medicine, 5(208), 208ra148–208ra148.

Mucha, P. J., Richardson, T., Macon, K., Porter, M. A., & Onnela, J.-P. (2010). Community structure in time-dependent, multiscale, and multiplex networks. Science, 328(5980), 876–878.

Newman, M. E. (2006). Modularity and community structure in networks. Proceedings of the National Academy of Sciences, 103(23), 8577–8582.

Nunes, C. S., Alonso, H., Castro, A., Amorim, P., & Mendonça, T. (2007). Towards the control of depth of anaesthesia: Identification of patient variability. In 2007 european control conference (ecc) (pp. 3109–3115).

Nunn, J. (1990). Effects of anaesthesia on respiration. BJA: British Journal of Anaesthesia, 65(1), 54–62.

Obert, D. P., Schweizer, C., Zinn, S., Kratzer, S., Hight, D., Sleigh, J., … Kreuzer, M. (2021). The influence of age on eeg-based anaesthesia indices. Journal of Clinical Anesthesia, 73, 110325.

Onoda, K., & Akama, H. (2023). Complex of global functional network as the core of consciousness. Neuroscience Research, 190, 67–77.

Pardo-Diaz, J., Bozhilova, L. V., Beguerisse-Díaz, M., Poole, P. S., Deane, C. M., & Reinert, G. (2021). Robust gene coexpression networks using signed distance correlation. Bioinformatics, 37(14), 1982–1989.

Parks, D. F., Schneider, A. M., Xu, Y., Brunwasser, S. J., Funderburk, S., Thurber, D., … Hengen, K. B. (2024). A nonoscillatory, millisecond-scale embedding of brain state provides insight into behavior. Nature Neuroscience, 27(9), 1829–1843.

Reisert, M., Mader, I., Anastasopoulos, C., Weigel, M., Schnell, S., & Kiselev, V. (2011). Global fiber reconstruction becomes practical. NeuroImage, 54(2), 955–962.

Romano, S., Vinh, N. X., Bailey, J., & Verspoor, K. (2016). Adjusting for chance clustering comparison measures. The Journal of Machine Learning Research, 17(1), 4635–4666.

Rubinov, M., & Sporns, O. (2010). Complex network measures of brain connectivity: uses and interpretations. NeuroImage, 52(3), 1059–1069.

Ruffini, G. (2017). An algorithmic information theory of consciousness. Neuroscience of Consciousness, 3(1).

Rukat, T., Baker, A., Quinn, A., & Woolrich, M. (2016). Resting state brain networks from eeg: Hidden markov states vs. classical microstates. arXiv preprint arXiv:1606.02344.

Sarasso, S., Casali, A. G., Casarotto, S., Rosanova, M., Sinigaglia, C., & Massimini, M. (2021). Consciousness and complexity: a consilience of evidence. Neuroscience of Consciousness, 2021(2), niab023.

Schwender, D., Kunze-Kronawitter, H., Dietrich, P., Klasing, S., Forst, H., & Madler, C. (1998). Conscious awareness during general anaesthesia: patients’ perceptions, emotions, cognition and reactions. British Journal of Anaesthesia, 80(2), 133–139.

Sen, B., Chu, S.-H., & Parhi, K. K. (2019). Ranking regions, edges and classifying tasks in functional brain graphs by sub-graph entropy. Scientific reports, 9(1), 7628.

Sevenius Nilsen, A., Juel, B. E., & Marshall, W. (2019). Evaluating approximations and heuristic measures of integrated information. Entropy, 21(5), 525.

Stevner, A., Vidaurre, D., Cabral, J., Rapuano, K., Nielsen, S. F. V., Tagliazucchi, E., … others (2019). Discovery of key whole-brain transitions and dynamics during human wakefulness and non-REM sleep. Nature Communications, 10(1), 1035.

Tan, X., Zhou, Z., Gao, J., Meng, F., Yu, Y., Zhang, J., … others (2019). Structural connectome alterations in patients with disorders of consciousness revealed by 7-tesla magnetic resonance imaging. NeuroImage: Clinical, 22, 101702.

Tasserie, J., Uhrig, L., Sitt, J. D., Manasova, D., Dupont, M., Dehaene, S., & Jarraya, B. (2022). Deep brain stimulation of the thalamus restores signatures of consciousness in a nonhuman primate model. Science advances, 8(11), eabl5547.

Terribilli, D., Schaufelberger, M. S., Duran, F. L., Zanetti, M. V., Curiati, P. K., Menezes, P. R., … Busatto, G. F. (2011). Age-related gray matter volume changes in the brain during non-elderly adulthood. Neurobiology of aging, 32(2), 354–368.

Urban, B., & Bleckwenn, M. (2002). Concepts and correlations relevant to general anaesthesia. British Journal of Anaesthesia, 89(1), 3–16.

Vidaurre, D., Quinn, A. J., Baker, A. P., Dupret, D., Tejero-Cantero, A., & Woolrich, M. W. (2016). Spectrally resolved fast transient brain states in electrophysiological data. NeuroImage, 126, 81–95.

Wang, R., Liu, M., Cheng, X., Wu, Y., Hildebrandt, A., & Zhou, C. (2021). Segregation, integration, and balance of large-scale resting brain networks configure different cognitive abilities. Proceedings of the National Academy of Sciences, 118(23), e2022288118.

Wang, Y., Ghumare, E., Vandenberghe, R., & Dupont, P. (2017). Comparison of different generalizations of clustering coefficient and local efficiency for weighted undirected graphs. Neural Computation, 29(2), 313–331.

Ward Jr, H. J. (1963). Hierarchical grouping to optimize an objective function. Journal of the American Statistical Association, 58(301), 236–244.

Warnaby, C. E., Sleigh, J. W., Hight, D., Jbabdi, S., & Tracey, I. (2017). Investigation of slow-wave activity saturation during surgical anesthesia reveals a signature of neural inertia in humans. Anesthesiology: The Journal of the American Society of Anesthesiologists, 127(4), 645–657.

Wilsenach, J. B. (2022). Computational network models for molecular, neuronal and brain data in the presence of long range dependence. *DPhil Thesis*.

Wilsenach, J. B., Warnaby, C. E., Deane, C. M., & Reinert, G. D. (2022). Ranking of communities in multiplex spatiotemporal models of brain dynamics. Applied network science, 7(1), 1–22.

Wu, L., Zhao, H., Weng, H., & Ma, D. (2019). Lasting effects of general anesthetics on the brain in the young and elderly:“mixed picture” of neurotoxicity, neuroprotection and cognitive impairment. Journal of anesthesia, 33, 321–335.

Yang, F. N., Picchioni, D., de Zwart, J. A., Wang, Y., van Gelderen, P., & Duyn, J. H. (2024). Reproducible, data-driven characterization of sleep based on brain dynamics and transitions from whole-night fmri. Elife, 13, RP98739.

Yeh, F.-C., Panesar, S., Fernandes, D., Meola, A., Yoshino, M., Fernandez-Miranda, J. C., … Verstynen, T. (2018). Population-averaged atlas of the macroscale human structural connectome and its network topology. NeuroImage, 178, 57–68.

Yeh, F.-C., Verstynen, T. D., Wang, Y., Fernández-Miranda, J. C., & Tseng, W.-Y. I. (2013). Deterministic diffusion fiber tracking improved by quantitative anisotropy. PloS one, 8(11), e80713.

Zamora-López, G., Zhou, C., & Kurths, J. (2010). Cortical hubs form a module for multisensory integration on top of the hierarchy of cortical networks. Frontiers in neuroinformatics, 4, 613.

Zhang, C., Bie, L., Han, S., Zhao, D., Li, P., Wang, X., … Guo, Y. (2024). Decoding consciousness from different time-scale spatiotemporal dynamics in resting-state electroencephalogram. Journal of Neurorestoratology, 12(1), 100095.

Zhang, Z., Cai, D.-C., Wang, Z., Zeljic, K., Wang, Z., & Wang, Y. (2019). Isoflurane-induced burst suppression increases intrinsic functional connectivity of the monkey brain. Frontiers in Neuroscience, 13, 296.

