## Supplementary Information for "Graph models of brain state in deep anaesthesia reveal sink state dynamics of reduced spatiotemporal complexity"

This section details supplementary tables and information for the study, organised into sections for each of the relevant methods used in the paper.

| subject ID | 1 | 2 | 3 | 4 | 5 | 6 | 7 | 8 | 9 | 10 | 11 | 12 | 13 |
|---|---|---|---|---|---|---|---|---|---|---|---|---|---|
| age | 21 | 24 | 21 | 23 | 30 | 29 | 29 | 33 | 19 | 25 | 32 | 25 | 34 |

*Supplementary Table 1:This table shows the anonymised subject ID and age of each participant used in the final analysis (after exclusion criteria were applied). The mean age of participants included in the study was 26.539±4.943, the median was 25, the minimum was 19 and maximum was 34.*

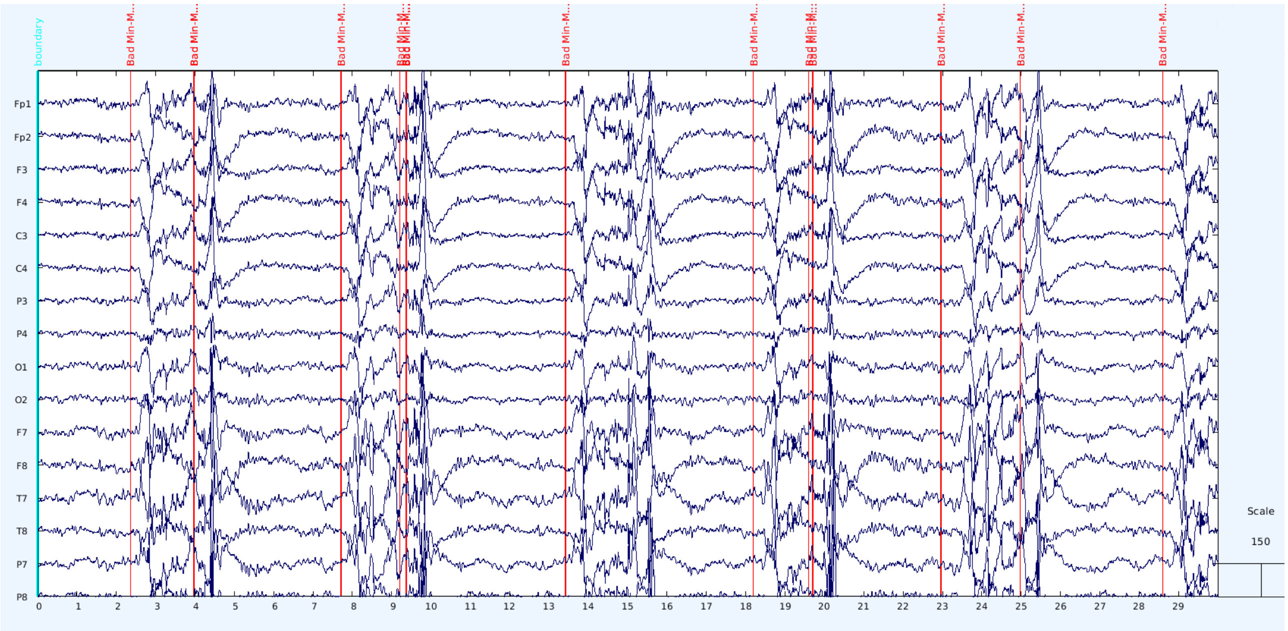


*Supplementary Figure 1: This figure shows the burst suppressed activity in a subset of electrodes recorded for one of the subjects that was excluded. The red Min-Max markers show the start location of local peaks in activity which are common in the burst regime of burst suppression. This figure was generated in EEGLAB (Makeig et al., 2004).*

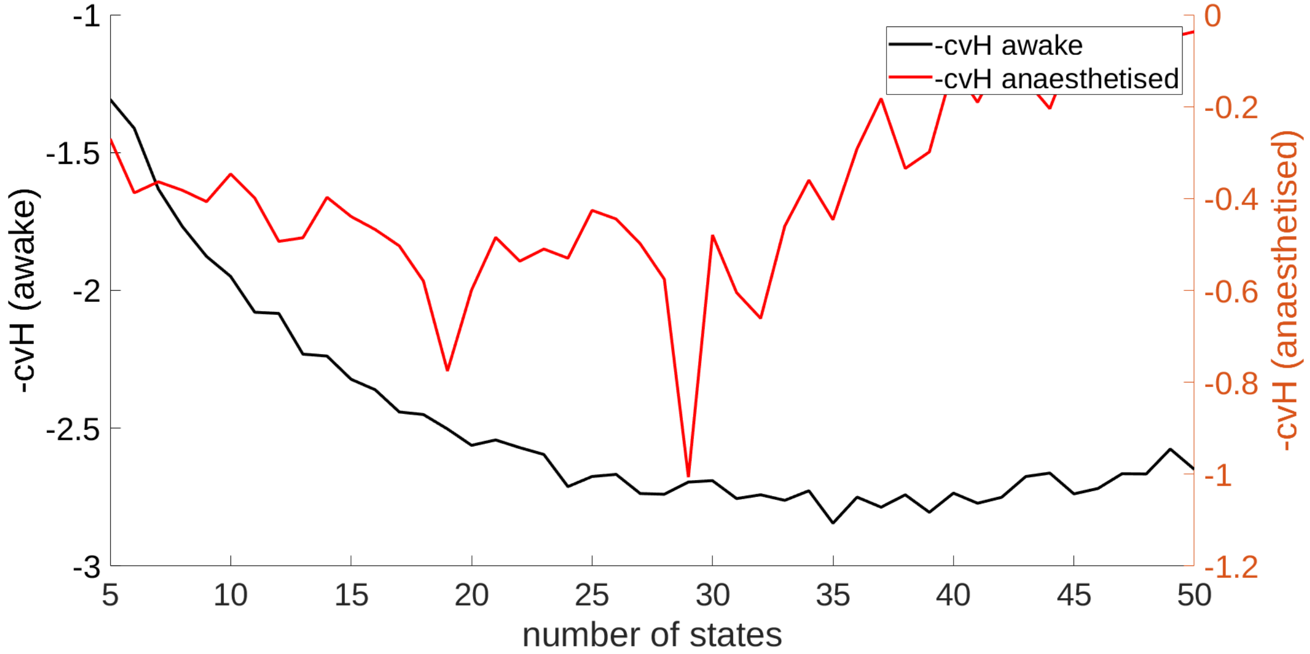


*Supplementary Figure 2: This figure shows the cross-validated (leave one subject out) entropy of the FO for both wakefulness (left) and anaesthesia (right) by which the initial number of total states was selected in both anaesthesia and wakefulness. The number of states tested ranges from* $K=5$ *to* $K=50$ *in both conditions. The method, first established in Wilsenach 2019, uses the first time the model approaches the local minimum of the cross-validated negative entropy (-cv-H). This was taken to be 33 for wakefulness and 19 for anaesthesia before removal of sporadic states to reach the final values of* $K_{wake}=32$ *and* $K_{anaes}=18$ *.*

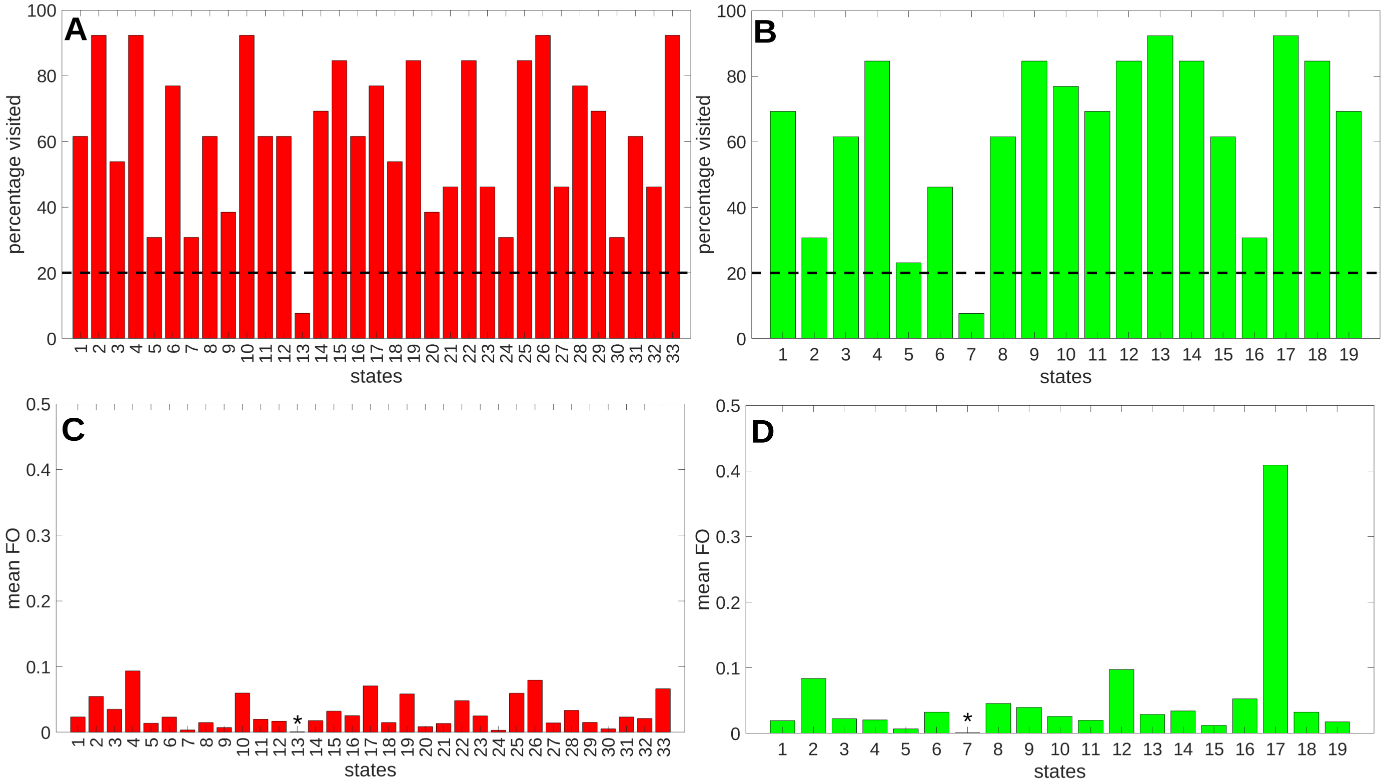


*Supplementary Figure 3: This figure shows the percentages of subjects visited in each state before the removal of rarely visited states in (A) wakefulness (with 33 initial states) and (B) anaesthesia (with 19 initial states). States were removed if they appear in less than 20% subjects shown as a dotted line. In addition, Fractional Occupancy (FO) of states are shown prior to removal in (A) wakefulness and (B) anaesthesia. A star (asterisk) is used to indicate the states which were removed in future analyses. It was observed that these states had an* $FO < 0.01$ *prior to removal.*

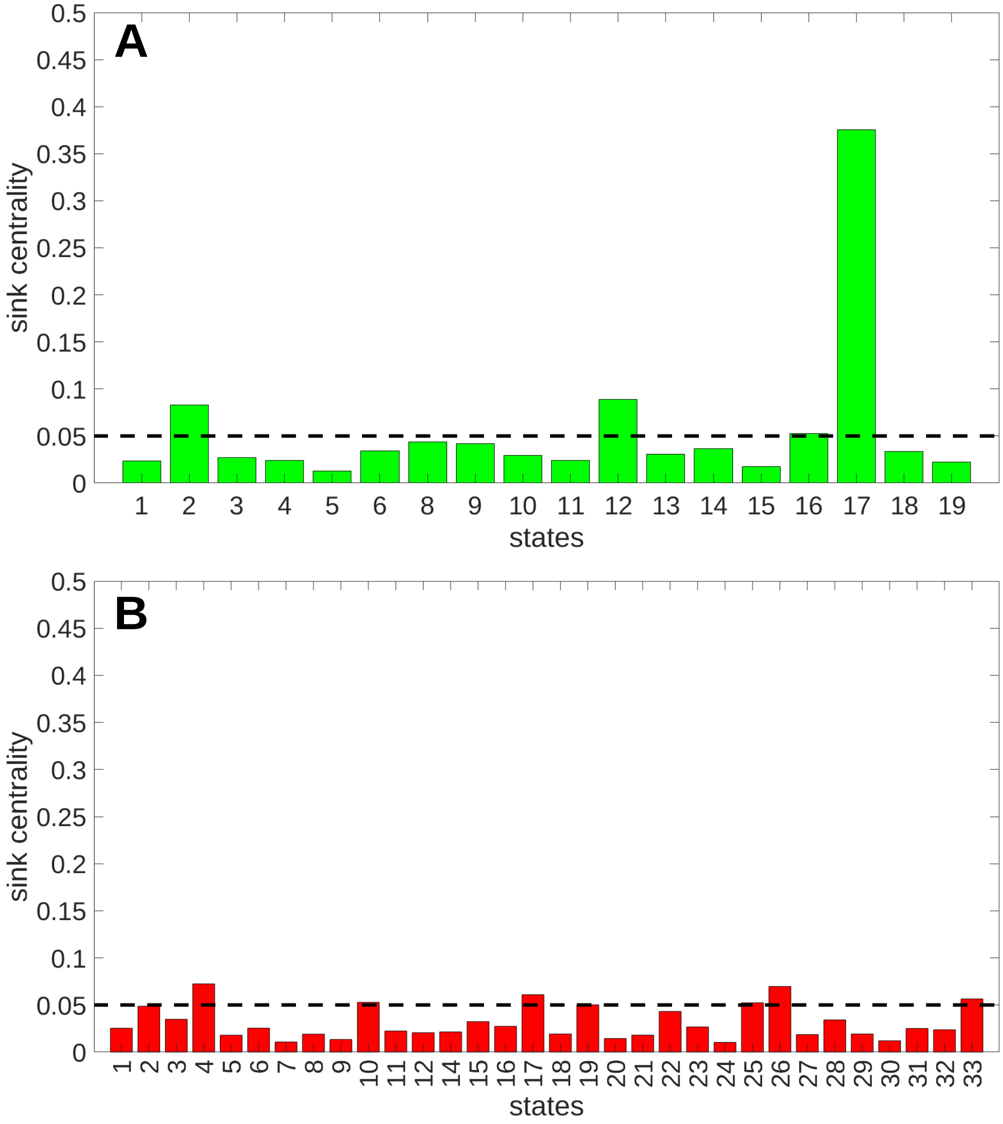


*Supplementary Figure 4: This figure shows the sink centrality of each state (stationary distribution over states) in (A) wakefulness and (B) anaesthesia. The strong sink state threshold of 0.05 is shown for both conditions as a dotted line.*

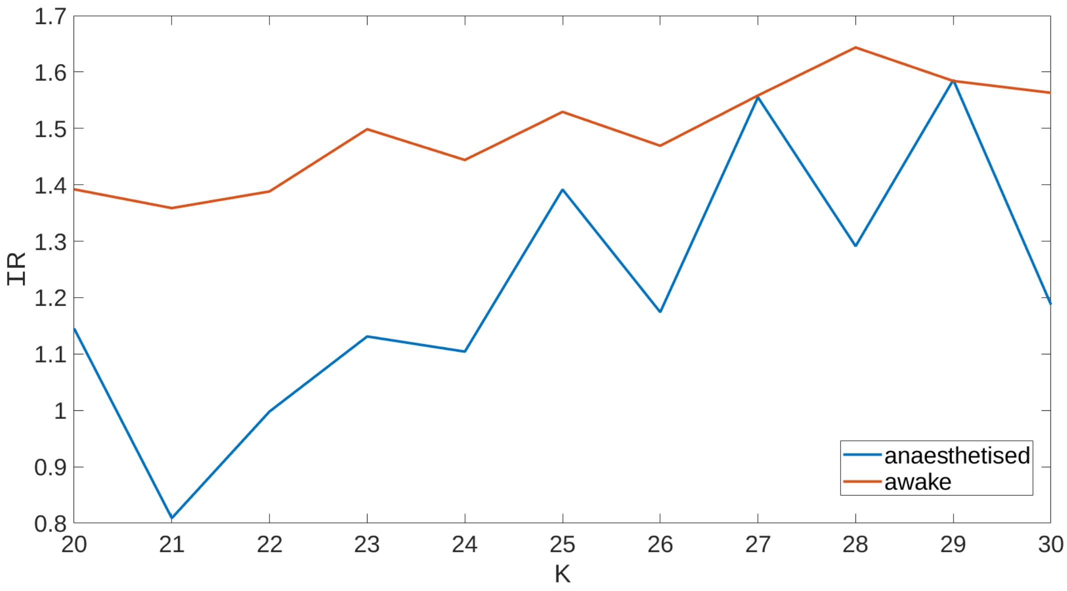


*Supplementary Figure 5: This figure shows the information rate (IR) in anaesthesia and wakefulness as a function of the number of states K. The information rate in anaesthesia is always at or below the information rate for wakefulness for all models having the same number of states*

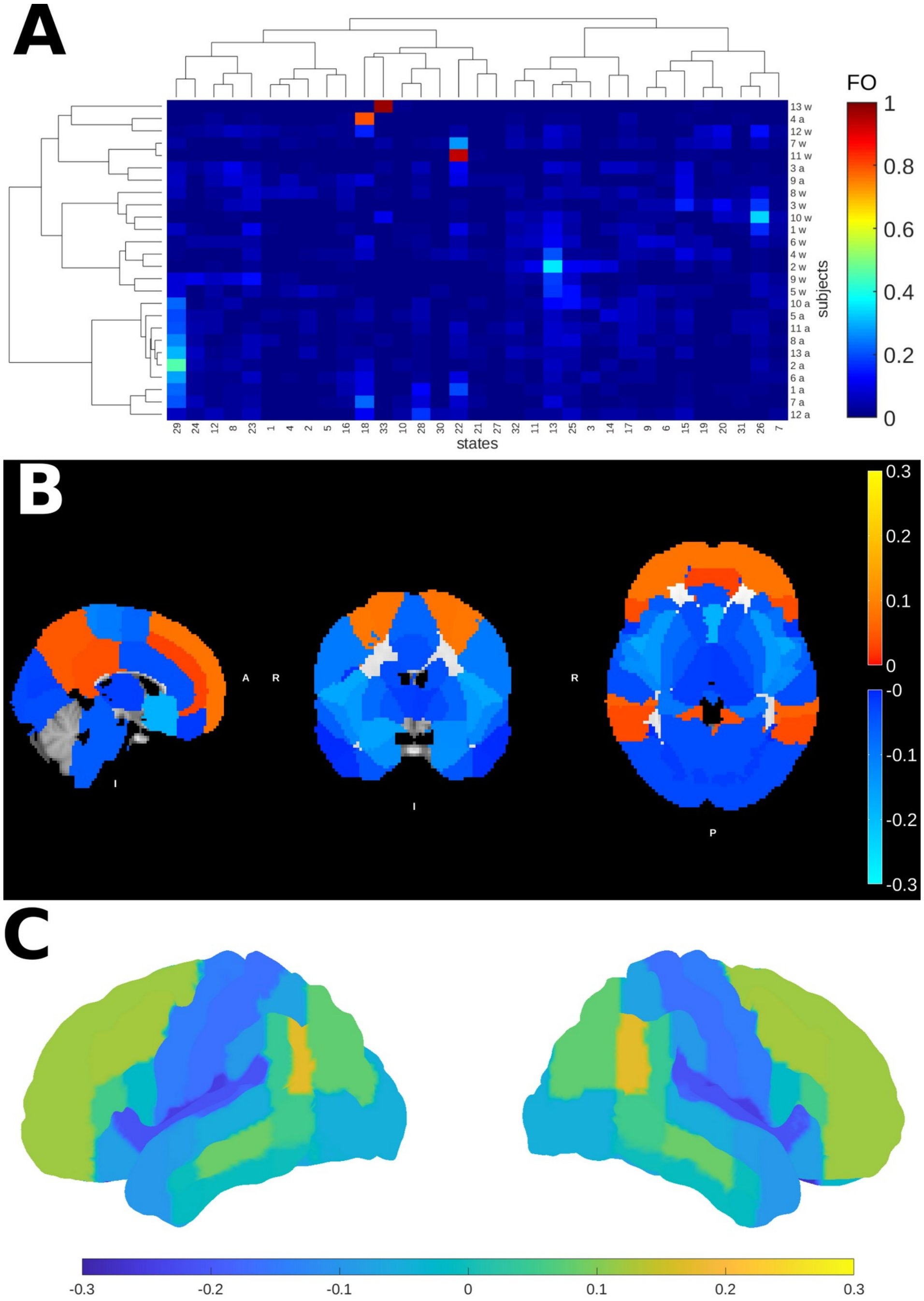


*Supplementary Figure 6: This figure summarises the results of training the HMGM on the combined anaesthesia-wakefulness trial data with* $K_{comb}=33$ *states (the same as the wakeful model).* ***(A)*** *This figure shows a clustergram of Fractional Occupancy (FO) over subject-trial with "w" referring to wakeful trials and "a" for deep anaesthesia trials. As in the original model, anaesthesia trials are dominated by a single state, state 29 in this case. As in the single condition anaesthesia model with* $K_{anaes}=18$ *, dominant sink state occurrence is positively correlated with age (* $p=0.005<0.01$ *, t-test, two-sided).* ***(B)*** *Cross-sectional activity maps of state 29 also show generally low absolute mean functional activity but exhibit clear increases in activity above baseline in the precuneous and prefrontal cortices. It is notable that the baseline here includes wakeful activity as well as deep anaesthesia activity and so cannot be readily compared to anaesthesia only model.* ***(C)*** *The surface plot of state 29 shows the extent of prefrontal activation in the dorsal prefrontal cortex relative to baseline.*

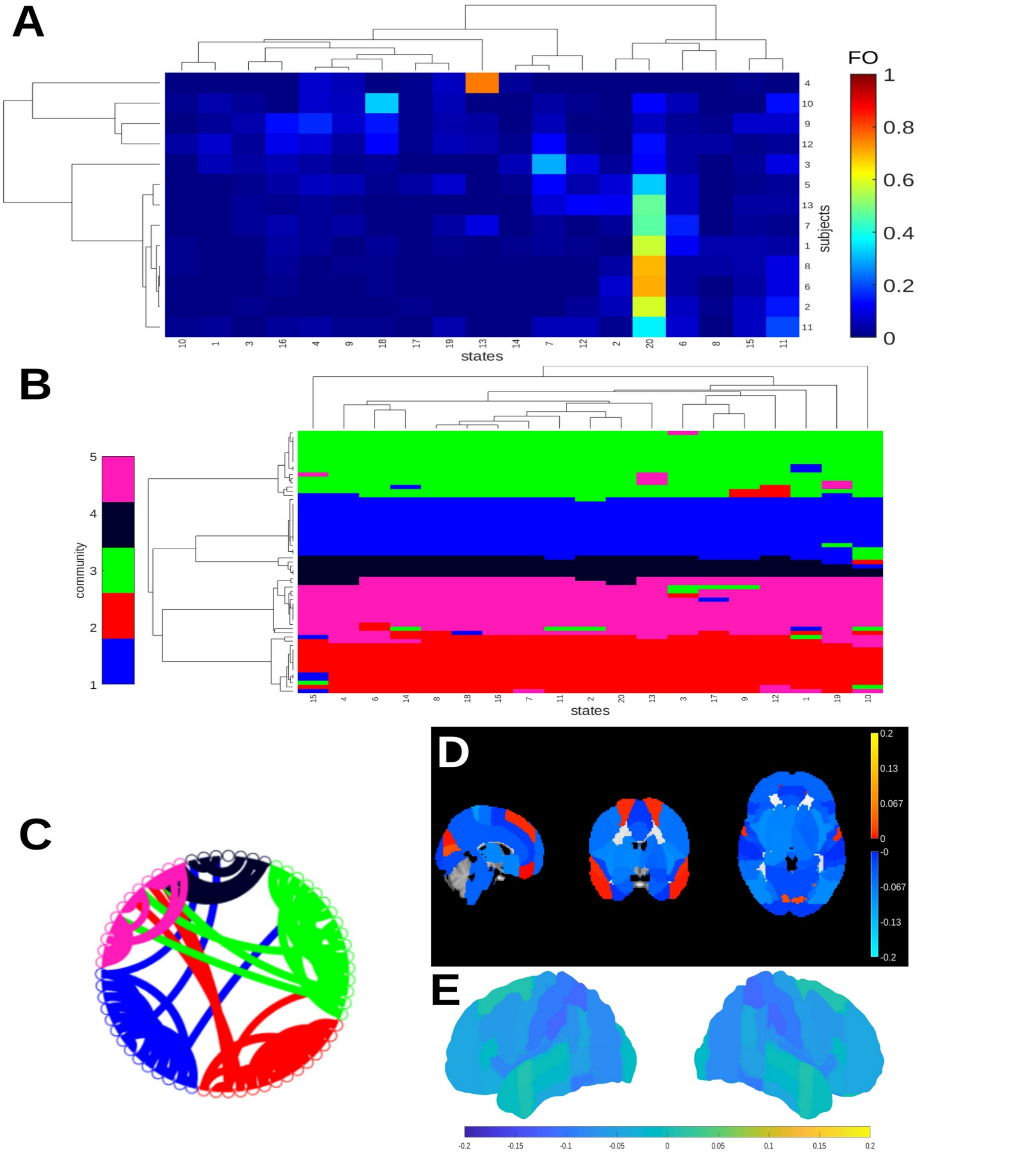


*Supplementary Figure 7: This figure summarises the results of fitting the anaesthesia model with one additional initial state, after state exclusion the resulting model has* $K_{anaes}=19$ *. states. (A) This figure shows a clustergram of Fractional Occupancy (FO) over subjects for the model. As in the* $K_{anaes}=18$ *anaesthesia model, subjects are dominated by a sink state, state 20. (B) The consensus community detection shows that stratification of activity in deep anaesthesia is conserved in the larger model, with the notable difference that most subcortical regions have combined into a single consensus community 1. (C) Sink state 20 connectivity is again relatively low between communities with low functional activity across the brain ((D) and (E)). These findings corroborate those for the simpler model with* $K_{anaes}=18$ *, and show that similar dominant sink states and dynamics emerge.*

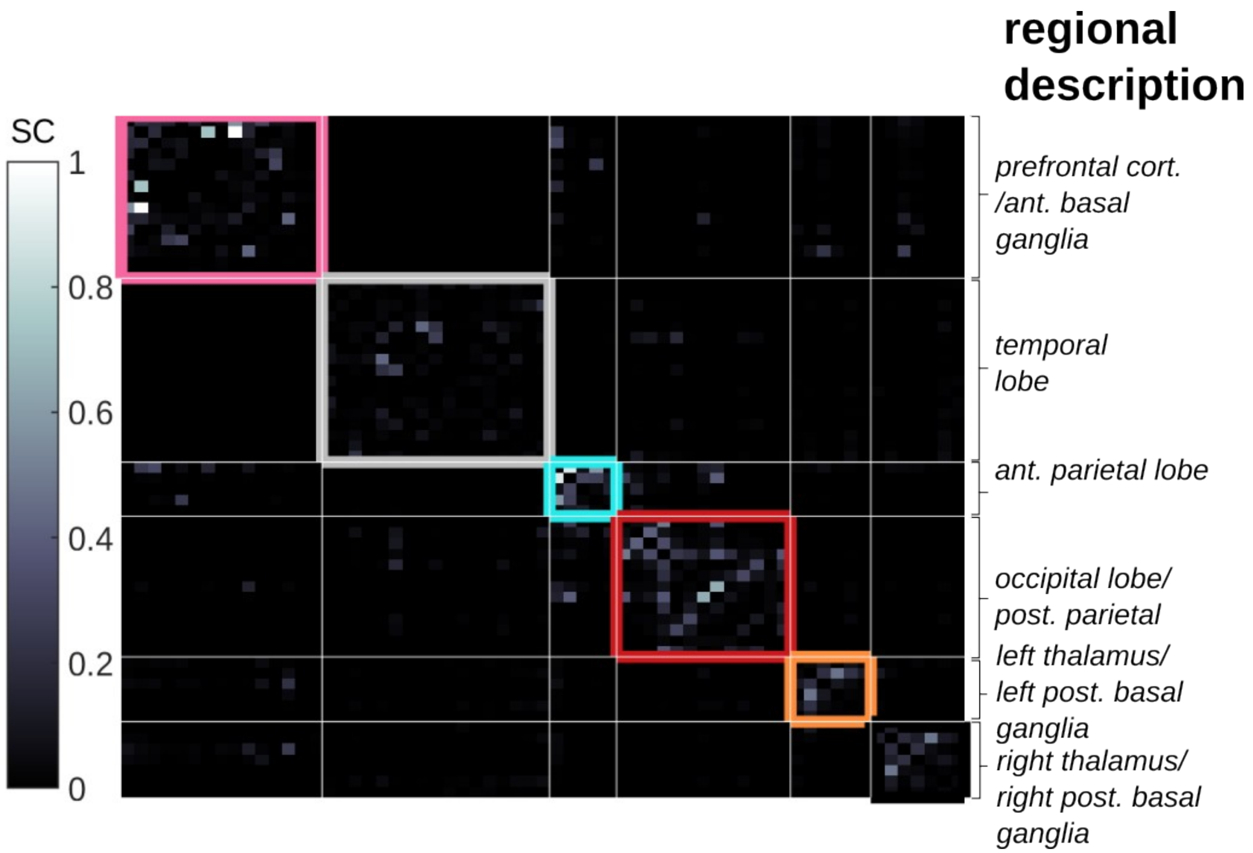


*Supplementary Figure 8: This figure summarises the structural connectivity and community composition of the structural graph model. Communities are shown as blocks with distinct colour borders and a general regional description is given to each community. These communities are the result of consensus community detection applied at the same resolution range as for the HMGM, where the range of values used is* $\Gamma=\{1,2,3\}$ *. Structural communities, like HMGM communities have a range of sizes and tend to be more lateralised than in the functional case due to the low levels of interhemispheric connectivity.*

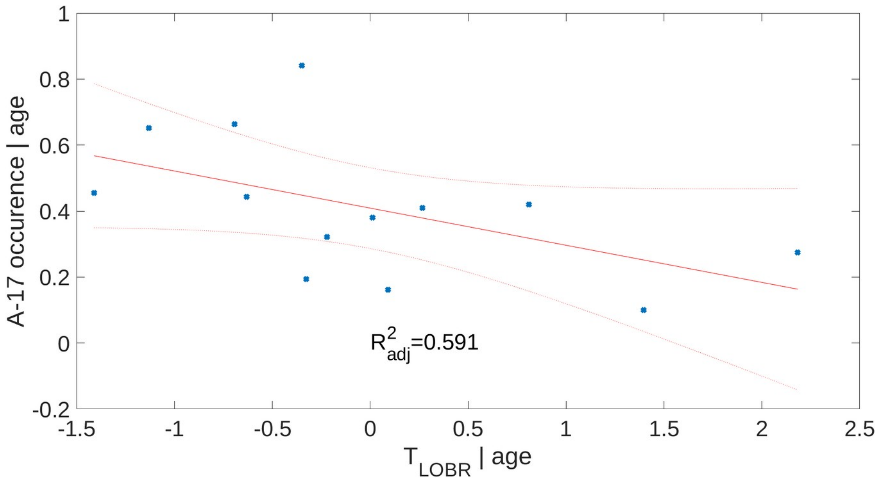


*Supplementary Figure 9: This figure shows the added variable plot for the multivariate linear regression of occurrence (in terms of FO) of sink state A-17, conditioned on age, against time at Loss of Behavioural Responsive ($T_{LOBR}$), also conditioned on age. The plot shows the residual data after fitting of the age only model (blue dots), the line of best fit (red line) has the slope equal to the effect size of $T_{LOBR}$ with 95% confidence interval (red dotted lines) shown ( $p=0.005$ , F-test). A significant positive relationship between A-17 occurrence and age persists ( $p=0.004$ , t-test, two-sided) and a marginally significant negative relationship between occurrence of A-17 and TLOBR is evident ( $p=0.079$ , t-test, two-sided). The model showed no significant deviation from normality of its residuals ( $p=0.981$ , Kolmogorov-Smirnov test) or heteroskedacity in its errors ( $p=0.131$ , Breusch–Pagan-Kroenker test).*

| variables | age | age+$T_{LOBR}$ | age+$GM_{LOBR}$ | age+$T_{LOBR}$+$GM_{LOBR}$ |
|---|---|---|---|---|
| BIC | 0.733 | -0.917 | 3.250 | 1.633 |
| $R_{adj}^2$ | 0.458 | 0.591 | 0.440 | 0.550 |

*Supplementary Table 2: This table shows adjusted $R^2$ ($R_{adj}^2$) the Bayesian Information Criterion (BIC), a Bayesian model selection criterion for the four models, shown in Supplementary Figures 2 and 3. For linear models, lower BIC is an indicator that a model better predicts the data given the number of dependent variables. Higher numbers of dependent variables are penalised by increasing the BIC. A Chi-squared was also performed to compare the likelihood of both models. The first model including only age was rejected in favour of the second model (age+$T_{LOBR}$),at the $p=0.040<0.05$ significance level (Chi-Squared likelihood ratio test).*

| variable | estimate | std. | t-statistic | p-value |
|---|---|---|---|---|
| intercept | -0.801 | 0.350 | -2.289 | 0.043 * |
| age | 0.046 | 0.013 | 3.513 | 0.005 ** |

*Supplementary Table 3: The results of fitting a multivariate linear model of A-17 occurrence (in terms of FO) against age. The table shows the estimated variable names, the estimated coefficients, the standard deviation of these coefficients, the t-statistics of each coefficient and the associated two-sided t-test p-value. The model explains the variance significantly better than the constant model at the $p=0.005<0.01$ significance level (F-test).*

| variable | estimate | std. | t-statistic | p-value |
|---|---|---|---|---|
| intercept | -0.34141 | 0.390 | -0.875 | 0.402 |
| age | 0.045574 | 0.013 | 3.713 | 0.004 ** |
| $T_{LOBR}$ | $3.365x10^{-4}$ | $1.719x10^{-4}$ | -1.957 | 0.079 (*) |

*Supplementary Table 4: The results of fitting a multivariate linear model of A-17 occurrence (in terms of FO) against age. The table shows the estimated variable names, the estimated coefficients, the standard deviation (std.) of these coefficients, the t-statistic of each coefficient and the associated two-sided p-value. The model explains the variance significantly better than the constant model at the $p=0.017<0.05$ significance level (F-test).*

| variable | estimate | std. | t-statistic | p-value |
|---|---|---|---|---|
| intercept | -0.302 | 2.617 | -0.116 | 0.910 |
| age | 0.045 | 0.014 | 3.256 | 0.009 ** |
| $GM_{cort.}$ | -0.883 | 4.591 | -0.192 | 0.851 |

*Supplementary Table 5: The results of fitting a multivariate linear model of A-17 occurrence (in terms of FO) against age and cortical grey matter density ($GM_{cort.}$). The table shows the estimated variable names, the estimated coefficients, the standard deviation (std.) of these coefficients, the t-statistic of each coefficient and the associated two-sided p-value. The model explains the variance significantly better than the constant model at the $p=0.023<0.05$ significance level (F-test).*

| variable | estimate | std. | t-statistic | p-value |
|---|---|---|---|---|
| intercept | -0.107 | 2.351 | -0.045 | 0.96475 |
| age | 0.043 | 0.012 | 3.444 | 0.007 ** |
| $T_{LOBR}$ | $-3.353x10^{-4}$ | $1.814x10^{-4}$ | -1.848 | 0.098 (*) |
| $GM_{cort.}$ | -0.418 | 4.127 | -0.101 | 0.921 |

*Supplementary Table 6: The results of fitting a multivariate linear model of A-17 occurrence (in terms of FO) against age. The table shows the estimated variable names, the estimated coefficients, the standard deviation (std.) of these coefficients, the t-statistic of each coefficient and the associated two-sided p-value. The model explains the variance significantly better than the constant model at the $p=0.017<0.05$ significance level (F-test).*

### S.1 Markov Models and Hidden Markov Models

The basic form of a finite state Markov model is as a dynamic model in which, for our purposes, a subject is said to occupy a state $s'$ , in a finite set $S$ of states, with some probability determined by the previous state $s$ . In the case that the model describes a stationary process, these probabilities can be summarised in terms of a matrix $P$ , with entries

$$P_{s,s'} = Pr(s \rightarrow s' | s) ,$$

where each entry describes the conditional probability of moving from state $s$ to $s'$ at some time given the current state $s$ .

A Hidden Markov Model (HMM) is one in which the state, $s$ is not directly observed but has to be inferred from the data. In our case we model each observation (time point) of the fMRI signal as arising from a multivariate Gaussian $O(s) \sim N(\mu(s), \Sigma(s))$ with mean and covariance dependent on the underlying state $s$ . Such a model is more difficult to fit than a model with fixed, state independent mean and covariance, but established approximate fitting procedures exist. Here, we use the minimal variational free energy approach applied in the HMMMAR package, a MATLAB package for learning of HMM models (Vidaurre et al. 2016). The model is trained using multivariate time series data $X$ , with consists of vector entries $X_t^n$ for $1 \leq t \leq T$ which are derived from the BOLD signal up to the final time $T$ and for each subject $n \leq N$ . As part of the fitting procedure, data was dimensionally reduced following the Parallel Analysis procedure used in[26] which uses randomised surrogate data to determine the optimal number of Principal Components to use in the reduction, see Supplementary Section 2.

First the eigenmatrix $U$ is calculated by Singular Value Decomposition (SVD) from the original dataset $X'$ , which has dimension $D \times TS$ , the reduced dimension $d < D$ is then calculated by Parallel Analysis (PA) and the final dataset $X$ is projected out using the reduced form of $U_d$ , with $d$ rows and used as input to train the HMM. The process is summarised below as

$$U_d X' = X .$$

After training, the number of states in each model is selected by maximising the entropy over the cross-validated maximum entropy of the Fractional Occupancy (FO), see Supplementary Figure 2. Fractional Occupancy (FO) is the fraction of time a subject $n$ is expected to spend in a state $s$ over a trial or recording, it is calculated from the a postereori probabilities of a state occurring across all time points in the recording. It can be represented in matrix form, with entries

$$FO_{n,s} = \sum_{1 \leq t \leq T} \frac{Pr(S_t = s | X)}{T} ,$$

where the $n^{th}$ row can be interpreted as a distribution representing the probability of observing subject $n$ across states in $S$ over the course of the recording. For more details on Markov models and the related HMMs see (Wilsenach 2022, Gagniuc 2017, Baum and Petri 1966). The motivation for this procedure is based on the statistical principle of maximum entropy which states that under conditions of only partial observations of the system (e.g. indirect observations of brain activity via the BOLD signal), the best model of the system is the model which retains maximal uncertainty about the unobserved components, while under the constraints of the model assumptions and available data (Jaynes, 1957).

### S.2 EEG-fMRI BOLD Acquisition and Experimental Set-up

In the original experiment, Blood oxygen level dependent (BOLD) fMRI images were obtained using a 3-Tesla human MRI system with a 1m bore magnet (Oxford Magnet Technology Ltd), a birdcage radio frequency coil and a reduced bore head gradient coil for pulse transmission (Magnex SGRAD MKIII. Magnex Scientific Ltd), and a four-channel helmet head coil for signal detection (PulseTeq). The system was controlled by a Varian Unity Inova console (Varian Medical Systems) using Siemens' gradients (Siemens Medical Ltd, Bracknell) (Mhuircheartaigh et al. 2013).

The experimenters used a whole brain gradient-echo echo-planar imaging sequence with acquisition of 41 contiguous coronal oblique slices of 3.5mm thickness inclined at 10 degrees to the axial plane (repetition time = 3000ms, echo time = 30ms, flip angle =90o, 224 x 224 mm field of view, 64 x 64 matrix). Four separate functional scans were performed to match the four phases of the experiment, i.e. resting state awake (10 minutes), induction (48 minutes), resting state – peak propofol dose (10 minutes) (Mhuircheartaigh et al. 2013).

Time at Loss of Behavioural Responsiveness ($T_{LOBR}$) was defined as the point during the induction phase where subjects failed to respond by pressing any of the buttons during a word identification task (regardless of correctness). The task was to identify whether consecutively played audio recordings of word pairs were the same or different, with two buttons corresponding to respective choices. Word pairs were presented at random with a mean interval of 56.8s and a range of 15s to 103.4s Mhuircheartaigh et al. 2013).

Two further scans were acquired at peak propofol dose while the subject was sedated. The first was a turbo flash T1-weighted (axial) high-resolution $1mm^3$ anatomical scan for co-registering the individual volunteer scans to standard stereotactic space (repetition time = 13ms, echo time = 5ms, flip angle $=8^o$, 256 x 192 mm field of view, 256 x 192 matrix). Finally a phase-difference image of B0, a fieldmap, was obtained for unwarping of functional scans (Mhuircheartaigh et al. 2013).

This protocol was repeated while recording from EEG only so as to obtain an EEG recording free of MRI-induced artifacts. This recording was used to determine subject exclusion on the basis of potential burst suppression at peak propofol dosages. EEG data for both experiments were acquired using a 32 channel EEG cap (BrainCap MR, Easycap GmbH) and MR compatible amplifier system (MRplus, BrainVision GmbH) at 5KHz sampling rate using FCz as a reference electrode. An abrasive electrolyte conducting gel was used between the electrodes and scalp to ensure electrode impedances were kept below 5kΩ. Filtering (high-pass=0.5 Hz, low-pass filter=70 Hz, notch=50Hz) was done with use of the acquisition software (BrainVision Recorder, version 1.10) that also recorded the timings of FMRI volumes (i.e. gradient artifact onset markers), stimuli presentation and subjects' button presses. Electrocardiographic signals and vertical/horizontal eye movements (VEOG/HEOG) were simultaneously recorded through an auxiliary device (BrainAmp ExG MR, Brain Products GmbH) for offline removal of the ballistocardiographic artifact and blink artifact removal, respectively (Mhuircheartaigh et al. 2013).

### S.3 Subject Exclusion Criteria

This study focuses on comparison with propofol-induced deep anaesthetic states while attempting to exclude subjects whose trajectory is dominated by atypical or potentially harmful activity. Burst-suppression is a state of deep anaesthesia that has been particularly associated with poor post-operative outcomes including post-operative delirium and cognitive decline (Fritz et al. 2016). It is characterised by a global pattern of alternating low and high amplitude periods of neural activity (see Supplementary Figure 1 for EEG activity example) (Zhang et al. 2019). We thus excluded subjects on the basis of pervasive burst-suppression (>2 minutes) during simultaneous EEG recordings. Recordings of subjects exhibiting brief periods of burst suppression (less than a minute across the recording) were not altered or removed. One subject was removed due to an incomplete anaesthesia recording, two subjects were removed on the basis of extensive burst suppression. This resulted in a reduced set of $N=13$ complete, deeply anaesthetised and awake subject recordings (see Supplementary Table 1) which were used in the construction of new wakefulness and anaesthesia HMGMs. The age of the final subjects ranged between 19 and 34 with a median of 25.

### S.4 Parallel Analysis for Dimensionality Reduction

Parallel analysis provides a principled approach to dimensionality reduction of the data $X'$ using SVD, resulting in the matrices $U_d$ above, as well as corresponding eigenvalues $\lambda_d$, for each dimension $d \leq D$. It is based on comparing the observed eigenvalues to a reference distribution. Since the reference (null) distribution of resting state fMRI BOLD signal eigenvalues is unknown we construct an empirical reference distribution based on independent shuffling of the fMRI BOLD signal across time and space in both wakefulness and anaesthesia data for each subject. This shuffling is repeated $R=10000$ times, producing $R$ surrogate data sets for each condition. PCA is then performed on each of these datasets producing a set of surrogate eigenvalues for each surrogate data set $\lambda_d^r$ for $r \leq R$. These values are then ordered from largest to smallest $\lambda_{(1)}^r \geq \ldots \geq \lambda_{(D)}^r$ and similarly for the original dataset $\lambda_{(1)} \geq \ldots \geq \lambda_{(D)}$.

The method works by defining a curve over dimensions $d$ determined by a percentile parameter $P$, such that $\lambda_{(d)}^P$ is the smallest possible value where $\lambda_{(d)}^r$ for $P$ % of the surrogate datasets. The optimal $d$ for dimensionality reduction is then defined as the minimum $d$ satisfying

$$\lambda_{(d+1)} < \lambda_{(d+1)}^P .$$

The parameter constrains the amount by which the eigenspectrum of the true data should dominate over the surrogate spectra, where in larger values for $P$ result in larger values for $d$. We choose a strenuous constraint of $P=99$ based on prior work on the fMRI BOLD signal in resting wakefulness and the recommendation that $P \geq 95$ in the literature (Humphreys et al. 1975, Wilsenach et al. 2022).

### S.5 Consensus Louvain Community Detection

Consensus community detection (as in the case of the structural connectome) was performed over a graph, $G=(V,A)$ , which is weighted and symmetric. This method improves on standard Louvain modularity-based community detection (Mucha et al. 2010), which uses the standard modularity score

$$Q(C)=\sum_{x,y\in V}\left[A^{x,y}-\gamma\frac{d^x d^y}{\sum_{u,v\in V}A^{u,v}}\right]\delta_C(x,y) ,$$

where $d^x$ is the weighted degree of $x\in V$ , $\gamma>0$ is the resolution parameter and

$$\delta_C(x,y)=\begin{cases}1 & \text{if } \exists c\in C \text{ such that } x,y\in c\\ 0 & \text{otherwise,}\end{cases} ,$$

is the Dirac delta for membership in the partition $C$ . The Louvain algorithm includes some randomness in partitioning and hence different runs may give different results. The robust method improves on directly optimising $Q(C)$ by considering a plausible range of resolution parameters $\{\gamma_1,\gamma_2,\ldots,\gamma_g\}=\Gamma$ and integrating the results of multiple realisations of the stochastic Louvain algorithm.

In consensus Louvain modularity optimisation the same modularity function for weighted, symmetric graphs, is optimised $R=1000$ times resulting in a partition of $V$ , denoted $C_{\gamma,r}$ for each repetition $1\le r\le R$ and $\gamma\in\Gamma$ . The consensus matrix

$$F_{x,y}=\frac{1}{R(\Gamma)}\sum_{\gamma\in\Gamma}\sum_{r=1}^{R}\delta_{C_{\gamma,r}}(x,y) .$$

is then formed, so that $F_{x,y}\in[0,1]$ . The final consensus community is obtained by performing Louvain community detection on the consensus matrix $F$ with resolution $\gamma=1$ . The resulting partition is generally robust to repeated applications of the Louvain algorithm. We select the values from $F_{s,s'}$ in increments of 0.1 in the interval $\Gamma=[0.1,1.5]$ . This range was chosen as for $\gamma>1.5$ , only the trivial partition (consisting of singletons) was produced for all transition graphs tested. This method is based on a simple method for consensus clustering (Lancichinetti and Fortunato 2012).

### S.6 Multiplex Graph Model and Spatiotemporal Consensus Community Detection

We introduce a novel approach to community detection that respects both spatial and temporal connectivity for dynamic state space models. Our new multi-resolution consensus community detection method is overall less vulnerable to the resolution limits of standard community detection methods and more robust to noise than conventional community detection approaches. The HMGM is a multiplex graph model with intralayer edges weighted by the transition probabilities in $P$ and interlayer edges determined by the functional connectivity $W(s)$ (analogous to $A$ in the structural case) between brain regions in state $s$ (Galatolo, 2010). The full spatiotemporal graph model is then $G(M)=(S\times V,A(W,P))$ with full spatiotemporal adjacency matrix

$$A(W,P)_{(s,x),(s',y)} = \begin{cases} P_{s,s'} & \text{if } x=y, s\neq s' \\ W(s)_{x,y} & \text{if } s=s' \\ 0 & \text{otherwise.} \end{cases} .$$

For a partition $C$ of $S\times V$ , the multiplex consensus community detection framework repeatedly optimizes the symmetric version of the multiplex modularity function

$$Q'(C) = \sum_{x,y\in V}\sum_{s,s'\in S}\left[W(s)_{x,y} - \gamma\frac{d(s)^x d(s)^y}{\sum_{u,v\in V} W(s)_{u,v}} + \omega P'_{s,s'}\right]\delta_C((x,s),(y,s'))$$

where

$$\delta_{C_{\gamma,\omega,r}}((s,x),(s',y)) = \begin{cases} 1 & \text{if } \exists C\in C_{\gamma,\omega,r} \text{ such that } (s,x),(s',y)\in C \\ 0 & \text{otherwise} \end{cases},$$

is the Dirac function, $P' = (P+P^T)/2$ , and $d(s)^x$ , $d(s)^y$ are the degrees of $x$ and $y$ in $W(s)$ , and $\omega$ is the temporal resolution which encourages shared community membership between regions in different states with shared modular organisation.

This definition of $Q'$ is a modified form of the standard multiplex modularity, which generally cannot take into account weighted or directed intralayer relationships. Models with constant temporal resolution values ( $\omega$ ) have been found not to be able to recover many simple multiplex patterns of community organisation (Hanteer and Magnani 2020). The addition of the non-constant inter-state transition probabilities $P'_{s,s'}$ allows for more complex interlayer relationships and thus potentially discovery of more complex spatiotemporal community structures. So far as we are aware, this is the first instance of leveraging the multiplex, temporal information encoded in multivariate HMMs to learn the repertoire of functional associations directly from recordings of functional brain activity. We used Louvain over the alternative Leiden optimisation algorithm due to the availability of the generalised algorithm for Louvain optimisation, which allows for such alternative formulations (Traag et al. 2019).

For the sake of robustness, as in the purely temporal case we perform consensus community detection. The relatively large multiplex community consensus matrix $F'$ combines community detection results from across temporal and spatial resolutions. Given $\Gamma' = \{\gamma_1,\ldots,\gamma_{g'}\}$ , $\Omega = \{\omega_1,\ldots,\omega_g\}$ and $R' = 1000$ replicates, $F'$ is a $DK\times DK$ matrix with entries given by

$$F'_{(s,x),(s',y)} = \frac{1}{R'|\Gamma||\Omega|}\sum_{\gamma\in\Gamma'}\sum_{\omega\in\Omega}\sum_{r=1}^{R'}\delta_{C_{\gamma,\omega,r}}((s,x),(s',y))$$

The resulting multiplex spatiotemporal consensus communities are then given by applying the standard Louvain algorithm to $F'$ with standard resolution $\gamma' = 1$ , resulting in the final partition $C'$ .

We chose the temporal resolution parameters $\Omega = \{0.01, 0.1, 1\}$ to maximise coverage of the temporal resolution space and spatial resolution $\Gamma' = \{1,2,3\}$, motivated by a previous study (Wilsenach et al. 2022), with $R' = 1000$.

### S.7 Stationary Distributions and The Information Rate

The stationary distribution $\pi_s$ of a transition probability matrix, $P$ of a regular Markov model (also known as a Markov chain), is the limiting long run probability of the occurrence of any one $s \in S$. It is defined as a vector over $S$ satisfying

$$\pi P = \pi .$$

A Markov model is said to be regular if the following positivity condition holds for some positive integer $m$, that is $P_{s,s'}^{(m)} > 0$ for all $s,s' \in S$, where $P^{(m)}$ is the result of taking $P$ to the power of $m$. In the case that the Markov model is regular, then there exists a unique stationary distribution with

$$IR(P) = \sum_{s,s' \in S} \pi_s P_{s,s'} \log(P_{s,s'}) ,$$

the information rate of $P$ (Cover, 1999). Here, we confirmed regularity for all transition matrices used in our models by validating the positivity condition for $m = 100$.

### S.8 Switching Rate and the Viterbi Path Inference for each Subject

The Viterbi algorithm is an established algorithm for estimating the most likely state trajectory (state sequence) given an HMM and a time series of observations for each subject under a condition,

$$S_n^* = \{ s_{n,1}^*, \dots, s_{n,T}^* \} .$$

Note that inference is performed independently for each trial meaning for each subject under each condition. The algorithm works in three steps: initialisation in which an initial probability is assigned to each state based on the observational model alone, induction in which the probabilities are updated using the probability of transition between states given the observations, and the termination where $S_n^*$ is computed by backtracking through the probabilities. For more details on this algorithm see (Viterbi, 1967).

Switching rates for each subject, $n$, are derived from the inferred trajectories by counting the state changes observed in $S_n^*$, $w_n^* = \#\{ s_{n,t}^* \neq s_{n,t+1}^* \}$.

### S.9 Efficiency Calculations and Normalisation

Global efficiency is a measure of the integration capacity of a given graph (or network) by virtue of its connections and the strength of those connections. Given the global efficiency $E_G$ of weighted network $G = (V, A)$ (with regions in $x,y \in V$ and weighted, symmetric adjacency $A$) as defined in (Rubinov and Sporns, 2010). It is defined in terms of the distance, $d_G(x, y)$, between regions in the graph which is determined from the inverse edge weights. The global efficiency is then,

$$EG_G = \frac{1}{N(N-1)} \sum_{x \neq y \in V} \frac{1}{d_G(x,y)},$$

where $N = |V|$. Similarly the local efficiency of a given node $x \in V$, $EL_G(x)$, is calculated by first constructing a graph $G^x$ from the edges between all neighbours of $x$, excluding $x$ itself. It is a measure of how well the local network stays integrated in the absence of $x$, or when $G$ is fully connected, how well the whole network is integrated in the absence of $x$. It is given as

$$EL_G(x) = \frac{\sum_{y,z \in V-\{x\}} \bar{A}^{x,y} \bar{A}^{x,z} \bar{d}_{G^x}(x,y)^{-1}}{\sum_{y,z \in V-\{x\}} \bar{A}^{x,y} \bar{A}^{x,z}}$$

where $\bar{d}_{G^x}(y,z)$ is the distance defined on $G^x$ when the edge weights, $A^{x,y}$, are replaced with $\bar{A}^{x,y} = (A^{x,y})^{1/3}$. This is the robust version of the local efficiency given in (Wang et al., 2016), which is modified from the original version proposed in (Rubinov and Sporns, 2010).

Efficiency measures need to be normalised by an appropriate null model. In order to control for the overall connectivity of the graph we use weight permutation as a null model against which to compare. Each null graph $G_r$ is produced by randomly permuting all entries in the upper triangular matrix based on $A$, $A^+$, with zeros everywhere else. Given such a permuted matrix, $A_r^+$, the new adjacency matrix of $G_r$ is defined,

$$A_r = \frac{A_r^+ + (A_r^+)^T}{2},$$

where $1 \leq r \leq R$. We then find the mean efficiency over all such $G_r$ and divide to obtain the connectivity normalised global efficiency,

$$EG_G^{'} = \frac{EG_G}{\frac{1}{N} \sum_{1 \leq r \leq R} EG_{G_r}}.$$

We set $R = 100$ here in order to generate a representative sample.

This measure was found to best match the degree distribution of the original graph when compared against the methods proposed in (Zalesky and Bullmore, 2012), for graphs based on correlation matrices, and (Rubinov and Sporns, 2011), for weighted graphs, using the Kolmogorov-Smirnov statistic to compare degree distributions across all brain state-related graphs in the study.

### S.10 Deriving Fractional Membership from Expected State Dynamics

Fractional membership brain maps were used to show the expected participation of each brain area in a given community in the long run average. These were derived from the following matrix defined over regions $x$ and communities $c \in C'$,

$$B_{c,x} = \sum_{s \in S} \pi_s \delta_c(s,x).$$

Here, $\delta_c(s,x)$ is the Dirac function that is one if and only if $x$ is in community $c$. This matrix therefore expresses a weighted average of the communities that $x$ is involved in given the time that the resting state brain is expected to spend in each state, wherein each row corresponds to a vector that can be mapped onto the parcellated brain maps shown in Figure 6 as fractional membership.

### S.11 Partition Similarity Measures

Two common comparison methods for partitions $C_1$ and $C_2$ of a vertex set $V$, are the Adjusted Mutual Information (AMI) and Adjusted Rand Index (ARI) (Romano et al. 2016). The two measures differ in their approach to determine partition similarity. The AMI is a measure of statistical dependence between the partitions, conceived of as a probability distribution over $V$. It is calculated as

$$AMI(C_1,C_2)=\frac{MI(C_1,C_2)-E_R\left[MI\left(C_1^R,C_2^R\right)\right]}{\max_{i\in\{1,2\}}\left\{H(C_i)\right\}-E_R\left[MI\left(C_1^R,C_2^R\right)\right]},$$

where $H(C_1)$ is the entropy of the partion, $MI(C_1,C_2)$ is the mutual information between partitions, $E_R[.]$ is the expectation assuming membership in both partitions were assigned uniformly at random and $C_i^R$ is a re-assignment of labels (repartitioning) of $C_i$.

On the other hand, the ARI quantifies the degree of shared membership between the partitions by counting the pairs of shared and unshared members across the partitions[76]. It adjusts for randomness according to some baseline model which is used to determine the expected amount of matching. Most commonly (and in our case), this is the permutation model which assumes that labels were randomly assigned. The ARI follows a similar form to the AMI by subtracting the expected value and dividing by the maximum

$$ARI(C_1,C_2)=\frac{RI(C_1,C_2)-E_R\left[RI\left(C_1^R,C_2^R\right)\right]}{\max_R\left\{RI\left(C_1^R,C_2^R\right)\right\}-E_P\left[MI\left(C_1^R,C_2^R\right)\right]}.$$

### S.12 Permutation Testing for Difference in AMI of Functional and Structural Communities

Permutation testing of which condition has the higher AMI when comparing communities to the structural models was carried out by permuting the columns of the two $S\times V$ community membership matrices $M_{wake}$ and $M_{anaes}$ of integer community labels. In this way, the distribution of communities across each state was preserved but the specific membership profile of each region is altered, resulting in surrogate membership matrices $M'_{wake}$ and $M'_{anaes}$. Each row (state) is then compared to the community structure of the structural model, producing surrogate $AMI'_{wake,i}$ and $AMI'_{anaes,j}$ for rows $i$ and $j$ respectively. The difference of the median AMI scores for both conditions surrogate AMI scores were then calculated. This was repeated $R=10^6$ times and the sample was compared to the observed difference in median AMI between models to produce a p-value based on the percentage of values that exceeded the observed score.

### S.13 Fractional Occupancy Permutation Testing

Permutation testing was performed by permuting the $N \times K$ FO matrix within each row, in order to produce surrogate FO distributions for each subject. The new matrices have the same size as the original matrix. This process is repeated $R = 10^5$ times to produce surrogate FO matrices. These new surrogate matrices remove any persistent across-subject structure of the FO matrix. We used these surrogates in order to test whether the observed correlation between sink state FO is significant. The two states with the highest mean FO across subjects were chosen for each subject FO matrices and the Pearson correlation between subject FO across states was calculated. The empirical p-value was then calculated from the percentage of surrogate correlations that exceed the actual value.

### S.14 Cortical Grey Matter Density Calculation

Cortical grey matter density was calculated from subject specific T1-weighted MRI images using the FSL-Fast package. FSL-Fast is a method which can be used to estimate fractional levels of occurrence of each of three tissue types in a T1 or high-resolution T2-weighted image: cerebral spinal fluid, white matter and grey matter. FSL-Fast uses a Hidden Markov random field approach to estimate voxel-wise fractional occurrences for each type and outputs the resulting brain maps for each type (Zhang et al. 2001). To calculate a subject-wise cortical grey matter density we first transformed the whole brain grey matter map to MNI152 standard space and then used a mask consisting of cortical only areas of the Harvard-Oxford cortical-subcortical parcellation to calculate mean grey matter density across the masked region.
